# Using empirical datasets to quantify uncertainty in inferences of landscape genetic resistance due to variation of individual-based genetic distance metrics

**DOI:** 10.1101/2021.03.01.432986

**Authors:** Joscha Beninde, Alain C. Frantz

## Abstract

1. Estimates of gene flow are commonly based on inferences of landscape resistance in ecological and evolutionary research and they frequently inform decision-making processes in conservation management. It is therefore imperative that inferences of a landscape factors relevance and its resistance are robust across approaches and reflect real-world gene flow instead of methodological artefacts. Here, we tested the impact of 160 different individual-based pairwise genetic metrics on consistency of landscape genetic inferences.
2. We used three empirical datasets that adopted individual-based sampling schemes and varied in scale (35-25,000 km^2^) and total number of samples (184-790) and comprise the wild boar, *Sus scrofa,* the red fox, *Vulpes vulpes* and the common wall lizard, *Podarcis muralis*. We made use of a machine-learning algorithm implemented in ResistanceGA to optimally fit resistances of landscape factors to genetic distance metrics and ranked their importance. Employed for nine landscape factors this resulted in 4,320 unique combinations of dataset, landscape factor and genetic distance metric, which provides the basis for quantifying uncertainty in inferences of landscape resistance.
3. Our results demonstrate that there are clear differences in Akaike information criteria (AICc)-based model support and marginal R^2^-based model fit between different genetic distance metrics. Metrics based on between 1-10 axes of eigenvector-based multivariate analyses (Factorial correspondence analysis, FCA; Principal component analysis, PCA) outperformed more widely used metrics, including the proportion of shared alleles (D_PS_), with AICc and marginal R^2^ values often an order of magnitude greater in the former. Across datasets, inferences of the directionality of a landscape factors influence on gene flow, e.g. facilitating or impeding it, changed across different genetic distance metrics. The directionality of the inferred resistance was largely consistent when considering metrics based on between 1-10 FCA/PCA axes.
4. Inferences of landscape genetic resistance need to be corroborated using calculations of multiple individual-based pairwise genetic distance metrics. Our results call for the adoption of eigenvector-based quantifications of pairwise genetic distances. Specifically, a preliminary step of analysis should be incorporated, which explores model ranks across genetic distance metrics derived from FCA and PCA, and, contrary to findings of a simulation study, we demonstrate that it suffices to quantify these distances spanning the first ten axes only.

## Introduction

Spatial genetic patterns form the basis for many ecological and evolutionary research avenues and are frequently utilized to guide decisions in conservation management. With the appropriate tools of analysis, patterns of isolation by distance (Wright (1943)), by resistance (McRae (2006)), by environment (Wang and Bradburd (2014)) or by barriers (IBB; Cushman, McKelvey, Hayden, and Schwartz (2006)) can be detected and reveal the extent to which a species’ environment can impact its gene flow pattern. Results of spatial genetic analyses are utilized in many different ways, e.g. to identify common drivers of patterns in the life-history of species (Medina, Cooke, & Ord, 2018), or to identify corridors and generate scientific guidance for conservation management, as is the goal, specifically, of many landscape genetic studies (Keller, Holderegger, van Strien, Maarten, & Bolliger, 2014). Underlying these efforts is the assumption that models of a landscapes’ impact on spatial genetic patterns are ecologically meaningful and robust to different methods of inference.

Landscape genetic approaches that statistically relate the distribution of genetic similarities among individuals to landscape characteristics (Cushman et al., 2006; Schwartz et al., 2009) hold great promise as several methodological problems have recently been solved. For example, linear-mixed effects models (LME) can account for non-independence of pairwise comparisons by the use of maximum-likelihood population effect (MLPE) parameterization (Shirk, Landguth, & Cushman, 2017b). Also, simulations have shown that LMEs with corrected Akaike information criteria (AICc) model-selection consistently outperform other regression methods, or Mantel-based methods, in correctly identifying true models of landscape impact (Shirk et al., 2017b). Furthermore, the recently-developed R-package ResistanceGA (Peterman, 2018), which combines LME models with MLPE parameterization, makes use of a machine-learning algorithm to infer resistance values of landscape factors that maximize fit to a user-specified genetic distance. This circumvents the inherent subjectivity and biases associated with user-specified, resistance values, which are frequently based solely on expert-opinion (Peterman et al., 2019; Peterman, Connette, Semlitsch, & Eggert, 2014; Richardson, Brady, Wang, & Spear, 2016).

However, approaches of landscape genetic inferences are still evolving (Balkenhol, Waits, & Dezzani, 2009; Manel & Holderegger, 2013; Richardson et al., 2016) and some methodological aspects remain underexplored. For instance, the genetic distance metrics used in (individual-based) landscape resistance modelling seems to be frequently chosen on *ad hoc* basis with no justification provided. The proportion of shared alleles, *D*_PS_, (Draheim, Moore, Fortin, & Scribner, 2018; E. L. Landguth, Cushman, Murphy, & Luikart, 2010; Trumbo, Spear, Baumsteiger, & Storfer, 2013) and other kinship- or relatedness-coefficients (Dellicour et al., 2019; Renner et al., 2016) are popular choices and continue to be implemented in new applications developed for landscape genetic analyses (Savary *et al.* 2020). However, many of these estimators have a large sample variance, which has recently been shown to negatively impact landscape genetic inferences (Winiarski, Peterman, & McGarigal, 2020). Also, Kimmig et al. (2020) have shown that AICc-based support and rank of the same single-surface resistance model can vary, depending on the genetic distance metric used for inference. These authors suggested that the use of a genetic distance based on ten axes of a factorial correspondence analysis (FCA; an eigenvector-based multivariate approach similar to principal component analysis, PCA) lead to model selection results with highest support (Kimmig et al., 2020). In contrast, results from a simulation study suggested that genetic distances derived from 64 PCA axes allows to correctly identify the underlying resistance surfaces most frequently (Shirk, Landguth, & Cushman, 2017a).

The uncertainty regarding the choice of genetic distance metrics and how it influences landscape genetic results led us to explore this issue more thoroughly. Basing ourselves on the ResistanceGA package and working with three distinct empirical datasets that employed individual-based sampling schemes, our overall objective was to test the impact of genetic distance metrics on model support and rank of different landscape factors and thereby to identify the most suitable genetic distance metrics for landscape genetic inferences. Expanding on a set of metrics used by Shirk et al. (2017a), we explored a total of 160 different individual-based genetic distance metrics. The majority of the additional metrics were derived from testing separately all PCA and FCA axes ranging from 1-75. Specifically, we aimed to test the prediction that, because they better emphasise differences between individuals, genetic distances based on eigenvector-based multivariate analyses consistently give rise to higher ranking models than the more commonly employed metrics that give all loci an equal weight. If this was the case, we wanted to assess whether one specific metric (FCA or PCA) in combination with a specific number of axes could be used to generalize recommendations for landscape genetic studies. Finally, we aimed to confirm that the resistance values inferred by the machine-learning algorithm of ResistanceGA are consistent for dataset-landscape factor combinations, independent of the genetic distance metric employed.

## Material & Methods

### Genotype datasets

The analyses were based on three previously published datasets: (1) The BOARS dataset consisted of 790 samples of a large omnivorous mammal, the wild boar (*Sus scrofa*) that were collected (2005-2009) randomly across the southern Walloon part of Belgium by the local Services of the Nature and Forest Department of the Public Service of Wallonia and genotyped at 14 microsatellite loci (Dellicour et al., 2019); (2) the FOXES dataset contained 184 samples of a medium-sized mammalian meso-predator, the red fox (*Vulpes vulpes*) that were road-killed (2010-2015) across the metropolitan area of Berlin and genotyped at 15 microsatellite loci (Kimmig et al., 2020); (3) the LIZARDS dataset contained 223 samples of a small insectivorous reptile, the common wall lizard, *Podarcis muralis* that were collected across the German city of Trier (2011-2012) and genotyped at 17 microsatellite loci (Beninde et al., 2016). The geographic distribution of samples of all three datasets is shown in **figure 1**. Wild boar and foxes are continuously distributed across the Walloon and Berlin study areas, respectively, while the lizard samples exhaustively cover all known distribution areas in Trier. The three datasets are thus especially suited to landscape genetic inferences, as they follow either a random sampling scheme in continuously distributed areas or cover all known occurrences. Such schemes are known to increase robustness of spatial genetic inferences and are preferred over clustered or transect sampling (Landguth & Schwartz, 2014; Schwartz & McKelvey, 2009).

**Figure 1:**
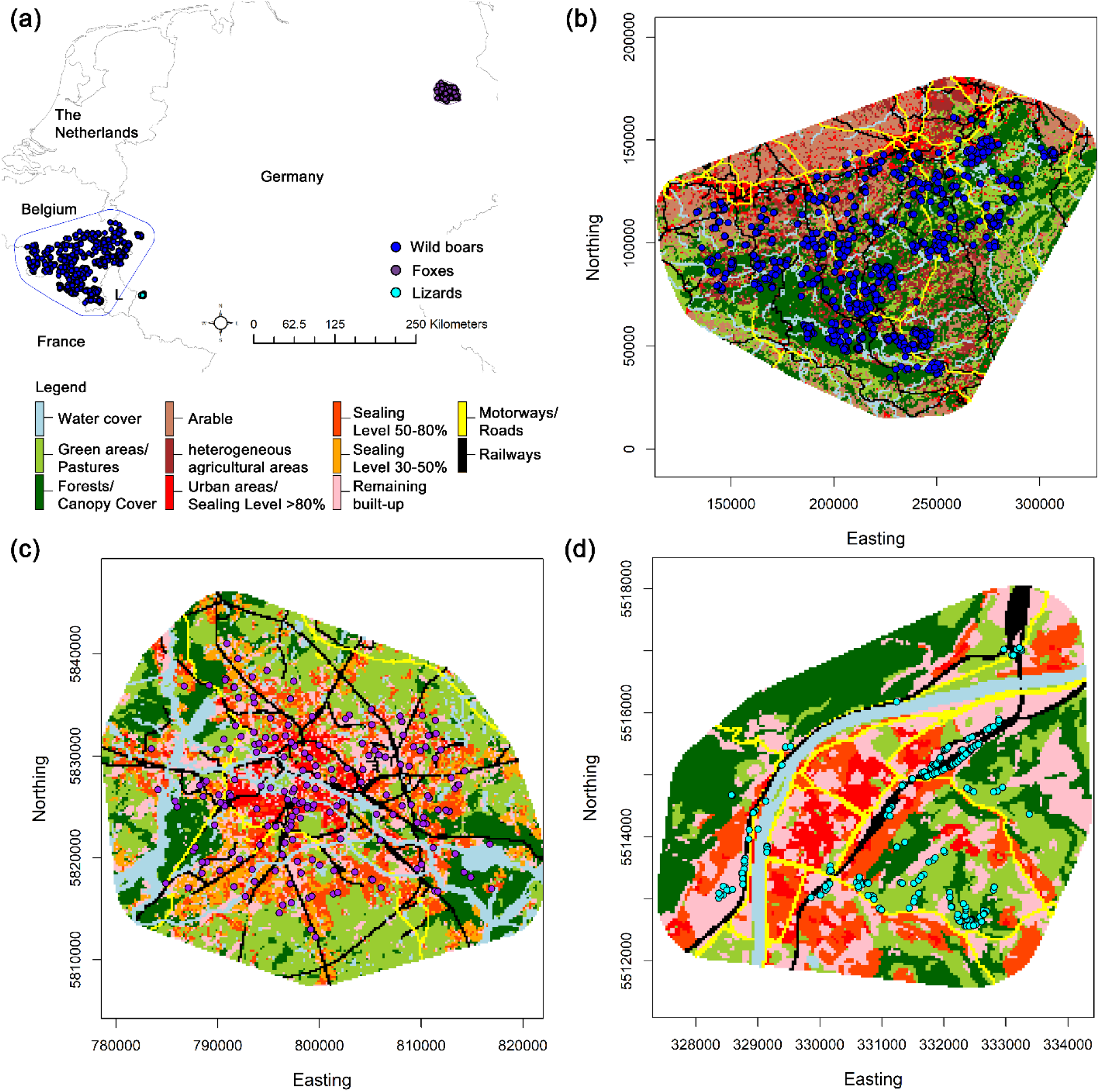
Geographic location of study regions and distribution of sample sites of all datasets shown jointly in Central Europe (a; L=Luxemburg). Land use types and samples sites (coloured dots) utilized in landscape genetic analyses are shown separately for BOARS (b), FOXES (c) and LIZARDS (d).

### Genetic distance metrics

For each of these three datasets, we computed a total of 160 different genetic distance metrics. Altogether 150 of these were based on two eigenvector-based multivariate analyses, the principal component analysis (PCA) and the closely related factorial correspondence analysis (FCA). Both approaches cluster variance between samples into composite gradients, accentuating differences between samples. PCA assumes continuous, normally distributed data (Dytham, 2011), whereas FCA assumes the data to consists of multistate categorical variables (She, Autem, Kotulas, Pasteur, & Bonhomme, 1987). In principle, the latter is thus more suitable for the analysis of microsatellite data. We used GENETIX v. 4.05.2 (Belkhir, Borsa, Chikhi, Raufaste, & Bonhomme, 1996-2004) to generate a contingency table of allele count (0, 1 or 2) by individual for all alleles in the population. We then used the R package ADE4 1.7.13 (Dray & Dufour, 2007) to perform FCAs and PCAs (with rescaling of allele counts) on the contingency table and obtain the position of every individual on every axis ranging from 1-75. For both FCA and PCA, we used the R package ECODIST 2.0.1 (Goslee & Urban, 2007) to obtain 75 genetic distances matrices based on the euclidean distance between positions of individuals on an incremental number of axes (first distance matrix based on the individuals’ position on axis 1, second distance based on positions on axes 1 & 2, etc.).

The remaining 10 genetic distance metrics represent other commonly used metrics and were also investigated by Shirk et al. (2017b). They can be grouped into the following categories. **Kinship coefficients:** (i) Kc.Lo (Loiselle, Sork, Nason, & Graham, 1995); (ii) Kc.R (Ritland, 1996); **relatedness coefficients:** (iii) Rc.L&R (Lynch & Ritland, 1999); (iv) Rc.Li (Li, Weeks, & Chakravarti, 1993); (v) Rc.Q&G (Queller & Goodnight, 1989); (vi) Rc.W (J. Wang, 2002); **fraternity coefficients:** (vii) Fc.L&R (Lynch & Ritland, 1999); (viii) Fc.W (J. Wang, 2002); **other metrics**: (ix) Rousset’s *â* (Rousset, 2000); (x) the proportion of shared alleles *D*_PS_ (Bowcock et al., 1994). For ease of reference, we will refer to these ten metrics as ‘Other Metrics’. The kinship, relatedness and fraternity coefficients are based on probabilities of alleles being identical by descent (using the sampled population as a reference). While the bias of these estimators tends to be small, they are generally characterized by a large sample variance (Lynch & Ritland, 1999; van de Casteele, Galbusera, & Matthysen, 2001; Vekemans & Hardy, 2004; J. Wang, 2002). Rousset’s *â* estimates genetic distance between individuals in a continuous population taking isolation-by-distance into account, while *D*_PS_ is a dissimilarity measure that considers the number of direct differences between genotypes. With the exception of *D*_PS_, which was estimated using the R package *adegenet* 2.1.1 (Jombart, 2008), all metrics were calculated using SPAGeDi v.1.5 (Hardy & Vekemans, 2002).

### Spatial datasets

For each genotype dataset we analysed nine landscape factors, resulting in a total of 16 factors (see t**able S1** for information on the source of the spatial data; **figures S1-S3** show all landscape model of BOARS, FOXES and LIZARDS, respectively), of which 15 landscape factors are depicted by binary surfaces while *slope* is depicted by a continuous surface. All landscape factors fall into the following three categories (see original publications for ecological justifications regarding choice of landscape factors; abbreviations correspond to datasets: B=BOARS; F=FOXES; L=LIZARDS). **Biotic factors**: (i) *green areas:* all types of arable land and grassland, fallow land, allotments, airports, public parks, cemeteries and bare soils [F,L]; (ii) *forests*: all forest irrespective of the composition [B,F]; (iii) *canopy cover*: all forested areas plus all street trees [L]; (iv) *hetero. agri*.: heterogeneous agricultural areas [B] (v) *arable*: arable land and permanent crops [B]; (vi) *pastures*: pastures [B]; **Anthropogenic infrastructure**: (vii) *motorways:* motorways only [B,F]; (viii) *roads*: all motorways, primary and secondary roads [L]; (ix) *railways:* all railway lines [B,L], including major stations [F]; (x) *urban areas:* all artificial surfaces [B]; (xi) *S.L. >80%:* sealing level > 80 %: continuous urban fabric [F,L]; (xii) *S.L. 50-80%:* sealing level 50 %-80 %: discontinuous dense urban fabric [F,L]; (xiii) *S.L. 30-50%:* sealing level 30 %-50 % discontinuous medium dense urban fabric [F]; (xiv) *remain. built-up:* the remaining Urban Atlas categories not covered by the previous categories [F,L]; **Abiotic factors**: (xv) *water cover:* water bodies, including major lakes, rivers and canals [B,F,L]; (xiv) *slope*: calculated from aggregated digital elevation models [B,L].

Proximity of a sampling location to the boundary of the study area can erroneously constrain the predicted movement of an individual during optimisation (Koen, Garroway, Wilson, Bowman, & Bersier, 2010). The extent of each study area was therefore defined by adding a buffer around a minimum convex polygon of the sampling locations whose width (BOARS: 20 km; FOXES: 5 km; LIZARDS: 1 km) reflected the upper margin of known, regular dispersal distances of the species in the specific area under investigation or in the type of habitat considered (Prévot & Licoppe, 2013; Schulte, 2008; Trewhella & Harris, 1990). Using ArcMap v.10.3 (ERSI Inc.), polygons were converted into rasters covering an area of 23,932 km^2^ with a resolution of 750 × 750 m in the case of the BOARS, of 1,206 km^2^ at 200 × 200 m resolution for FOXES and of 34 km^2^ at 40 × 40 m resolution for LIZARDS. We increased grid cell size for FOXES and LIZARDS compared to the original publications in order to increase computational speed. For each study area we generated a single polygon containing all environmental features of interest (except *slope*) and converted this polygon into a raster by categorically classifying each cell as the single feature with the largest area within the cell. However, every ‘linear’ feature (*motorways*, *roads*, *railways*, *water cover*) in the single input polygon was given a positive priority value. Thus, when a grid cell overlapped with a linear feature, it was categorically assigned to that feature, irrespective of the proportion of the cell this feature occupied. However, each linear feature was given the same priority value so that when two or more linear features overlapped with a grid cell, the cell was classified as the linear feature that occupied the greatest proportion of the cell. The ensuing raster with all the categorical features was re-classified into eight or nine single-feature rasters (depending on whether *slope* was considered) where grid cells had a value of one or zero, depending on whether it contained the feature of interest.

### Landscape genetic analyses

Optimizations of resistance values were conducted using ResistanceGA 4.0.14. As mentioned above, this R package makes use of a machine-learning algorithm to infer resistance values of landscape factors and combines LME models with MLPE parameterization and AICc-based model-selection. We used the *ss_optim*() function to optimize the resistance of single environmental predictors and test model selection performance of the genetic distance metrics. We used the *commuteDistance* function of the R-package gdistance v.1.2-1 (van Etten, 2017) to calculate pairwise resistance distances between individuals, which is equivalent to circuit-theory-based resistance distances (Kivimäki, Shimbo, & Saerens, 2014). Using the *GA.PREP*() function, we defined a range of 1-500 for the resistance values of categorical surfaces to be assessed during optimization. We refrained from expanding the explored parameter space when optimised resistance values were at or near the 500 limit, as we did not aim to identify precise resistance values, but rather aimed to quantify variation of landscape genetic inferences across genetic distance metrics. For unknown reasons we obtained resistance estimates >500 for one landscape factor in the BOARS dataset and for five landscape factors in the FOXES dataset (see Results).

Model fit was evaluated based on AICc: if the difference in AICc (ΔAICc) between two models was >2 AICc units, the model with the smallest AICc value was considered to be a better fit. For each optimized resistance surface, we calculated the difference between its AICc value and that of the corresponding *distance-only* model (we will refer to this measure as Δdist-AICc) and used this ‘distance-corrected’ AICc value for comparisons of model performance across genetic distance metrics and datasets. The model with the largest (positive) Δdist-AICc value was considered to be the best-supported mode. If the Δdist-AICc of an individual predictor was <2 AICc units, we considered this specific model not to be better supported than the *distance-only* model.

Optimising each one of the nine single-feature surfaces per dataset separately using 160 different genetic distance metrics amounted to a total of 4,320 unique combinations of genotype dataset, landscape factor and genetic distance metric. To account for stochastic parameter estimation of the machine-learning algorithm, all unique combinations we optimized twice (resulting in a total of 8,640 optimizations). However, we considered only the higher-ranking model of each paired optimisation run when next performing a (pseudo-)bootstrap procedure with the *resist.boot*() command. This function sub-samples a user-specified proportion of the individuals without replacement, refits the MLPE model based on the already optimised resistance surface and re-calculates AICc values. This procedure should provide for robust model selection, independently of the combinations of samples in the input file. We performed the bootstrap procedure simultaneously for all resistance surfaces of a specific combination of genotype dataset and genetic distance metric, sampling 75% of the observations at each of 1000 iterations, and compared the relative fit of these models across different genetic distance metrics based on Δdist-AICc values.

The optimised single-feature resistance surface consists of a binary grid where cells classified as the factor under investigation all have the same value and all the other cells have another value. If the optimisation process infers that a landscape feature under investigation resists gene flow, it is given a value ranging from >1 to 500 (as specified above), while all the other cells are given a value of 1. In the opposite case, when a landscape feature facilitates gene flow, it is given a value of 1 and all the other cells receive values ranging from >1 to 500, i.e. indicating a resistance to gene flow. For easier comparison, we combined resistance values of both scenarios by expanding the range of resistance scores from −500 to 500. In the case of a landscape factor resisting gene flow, we retained the resistance values, while in the case of a feature inferred to conduct gene flow, we replaced its resistance value of 1 with that of the other cells, changed to a negative number. Therefore, based in this combined resistance measure, if resistance values of a single feature was −500, it is a strong conductor of gene flow, while a resistance score of 500 indicates strong resistance to gene flow. We also retained larger resistance values, in the few cases where we obtained resistance estimates >500.

## Results

### Bootstrapping and creating final dataset

The difference in AICc values between the models of two runs of optimization for identical dataset-landscape factor-genetic distance metric combinations was <1 AICc in 3631/4,320 cases (84 %), while the difference ranged from 1 to 303 in the remaining 689 combinations (median = 8.56; see histogram in **figure S4**). For every identical combination, we used the model of the optimisation run with higher AICc value for bootstrapping and all further analyses (for Δdist-AICc values of both optimization runs of FCA and PCA see **figures S5** for BOARS, **S6** for FOXES and **S7** for LIZARDS). The differences in Δdist-AICc values between the initial runs of optimization and bootstrapping results were mostly small (correlation coefficient = 0.994; **figure S8**). Even though the range of resistance values to be explored during optimisation was set to 1-500 via the GA.PREP() function, for unknown reasons we obtained resistance estimates beyond this parameter space for one landscape factor in BOARS and for five landscape factors in FOXES. Models that had a wider parameter space disposable for optimization had significantly higher AICc values when compared to models of identical dataset-landscape factor-genetic distance metric combinations that stayed within the set range (Wilcoxon signed rank test, V=464, P<0.001), with a median difference of 4.55 Δdist-AICc (ranging from 0.1-62.9 Δdist-AICc; see histogram in **figure S9**). Since this difference in Δdist-AICc values was small in comparison to differences between ranks in the final table of results (**table 1 & S2**) and as it was spread across the majority of genetic distance metrics (151 out of 160), we included all results in subsequent analyses.

**Table 1:** Results of genetic distance metrics derived from the first ten axes of FCA and PCA and all Other metrics.

| *Genetic distance metric* | *BOARS* |  |  | *FOXES* |  |  | *LIZARDS* |  |  |
| --- | --- | --- | --- | --- | --- | --- | --- | --- | --- |
| | *Landscape factor* | $\Delta dist$<br>*AICc* | $\Delta dist$<br>*AICc*<br>>2 | *Landscape factor* | $\Delta dist$<br>*AICc* | $\Delta dist$<br>*AICc*<br>>2 | *Landscape factor* | $\Delta dist$<br>*AICc* | $\Delta dist$<br>*AICc*<br>>2 |
| **DPS** | urban areas | 20.95 | 7 | motorways | 18.35 | 2 | S.L. 50-80% | 7.75 | 2 |
| **Fc.L&R** | arable | 9.38 | 1 | railways | 1.50 | 0 | water cover | 86.97 | 4 |
| **Fc.W** | railways | 5.85 | 2 | water cover | 3.03 | 2 | S.L. >80% | -15.37 | 0 |
| **Kc.Lo** | arable | 81.23 | 7 | forests | 10.33 | 4 | S.L. 50-80% | 1.59 | 0 |
| **Kc.R** | arable | 105.9<br>4 | 8 | forests | 7.77 | 4 | slope | 1.98 | 0 |
| **Rc.L&R** | arable | 92.18 | 8 | forests | 10.63 | 4 | S.L. >80% | 7.71 | 3 |
| **Rc.Li** | water cover | 29.40 | 7 | motorways | 20.08 | 3 | S.L. 50-80% | 2.41 | 1 |
| **Rc.Q&G** | pastures | 23.17 | 4 | motorways | 14.75 | 2 | water cover | 24.69 | 7 |
| **Rc.W** | water cover | 31.45 | 7 | motorways | 16.92 | 2 | roads | 1.77 | 0 |
| **Rousset's a** | pastures | 51.24 | 9 | motorways | 21.23 | 3 | water cover | 660.60 | 8 |
| **FCA_1** | forests | 477.5<br>2 | 9 | water cover | 212.5<br>7 | 8 | roads | 26.66 | 7 |
| **FCA_2** | railways | 1427.<br>20 | 8 | water cover | 304.0<br>9 | 9 | S.L. 50-80% | 25.30 | 4 |
| **FCA_3** | motorways | 761.4<br>9 | 8 | water cover | 277.9<br>2 | 9 | S.L. 50-80% | 24.83 | 7 |
| **FCA_4** | motorways | 673.8<br>1 | 8 | water cover | 134.2<br>0 | 9 | slope | 38.83 | 8 |
| **FCA_5** | motorways | 346.7<br>2 | 9 | water cover | 89.41 | 9 | water cover | 2268.7<br>9 | 9 |
| **FCA_6** | motorways | 408.4<br>9 | 9 | water cover | 75.42 | 8 | water cover | 2416.4<br>1 | 9 |
| **FCA_7** | motorways | 333.7<br>9 | 8 | water<br>cover | 80.51 | 8 | water cover | 1433.4<br>2 | 9 |
| **FCA_8** | motorways | 286.3<br>1 | 8 | water<br>cover | 78.20 | 8 | water cover | 995.34 | 9 |
| **FCA_9** | motorways | 270.2<br>1 | 8 | water<br>cover | 79.92 | 9 | water cover | 928.54 | 9 |
| **FCA_10** | motorways | 225.4<br>3 | 8 | water<br>cover | 70.61 | 8 | water cover | 815.14 | 9 |
| **PCA_1** | forests | 994.6<br>3 | 8 | water<br>cover | 208.5<br>2 | 8 | roads | 839.83 | 8 |
| **PCA_2** | railways | 555.9<br>9 | 8 | water<br>cover | 83.96 | 9 | water cover | 2111.0<br>3 | 9 |
| **PCA_3** | motorways | 193.6<br>5 | 8 | water<br>cover | 56.73 | 8 | water cover | 2279.9<br>9 | 9 |
| **PCA_4** | forests | 192.5<br>3 | 8 | water<br>cover | 52.07 | 9 | water cover | 2017.1<br>5 | 9 |
| **PCA_5** | forests | 139.1<br>7 | 8 | water<br>cover | 59.40 | 8 | water cover | 2104.1<br>6 | 9 |
| **PCA_6** | forests | 131.1<br>7 | 8 | water<br>cover | 42.55 | 8 | water cover | 1677.8<br>9 | 9 |
| **PCA_7** | motorways | 122.8<br>4 | 8 | water<br>cover | 37.52 | 8 | water cover | 1183.8<br>9 | 9 |
| **PCA_8** | motorways | 123.5<br>0 | 8 | water<br>cover | 42.91 | 8 | water cover | 951.61 | 9 |
| **PCA_9** | motorways | 109.0<br>8 | 8 | water<br>cover | 40.21 | 8 | water cover | 733.32 | 9 |
| **PCA_10** | motorways | 136.5<br>9 | 8 | water<br>cover | 38.26 | 9 | water cover | 630.28 | 9 |

### Influence of genetic distance metrics on model ranks, Δdist-AICc value and marginal R^2^

AICc-based model rank of all 27 landscape factors analysed across the three datasets varied depending on the choice of genetic distance metric used for analysis (**figure 2**). Depending on the distance metric employed, seven different landscape factors received highest AICc-based model support in BOARS and five different landscape factors in the case of both the FOXES and LIZARDS datasets. There was a significant effect of genetic distance metric on the median Δdist-AICc values of all nine single-feature models in each dataset (BOARS: Kruskal-Wallis χ^2^ = 258.8, d.f.=159, P<0.001; FOXES: χ^2^ = 198.2, d.f.=159, P=0.019; LIZARDS: χ^2^ = 446.3, d.f.=159, P=0.001; **figure 3**), with the first axes of the FCA- and PCA-derived genetic distance metrics consistently yielding the highest median Δdist-AICc values (**figure 3**). For each landscape factor bar one, the highest Δdist-AICc values were obtained by models using FCA- or PCA-derived genetic distance metrics based on between 1-10 axes. These were frequently up to an order of magnitude greater than the highest Δdist-AICc values inferred using models (**table 1 & S2**). Consequently, pairwise comparisons showed that the Δdist-AICc values were significantly higher when employing FCA- or PCA-derived genetic distance metrics based on between 1-10 axes, then when using genetic distance metrics based on all ten Other Metrics or on FCA and PCA axes 11-75 (**table 2; figures S10-S12**). Further comparing Δdist-AICc values between models based on axes 1-10 of an FCA with axes 1-10 of a PCA yielded variable results: FCA 1-10 gave rise to significantly higher Δdist-AICc values in BOARS and higher, marginally insignificant Δdist-AICc values in FOXES, while PCA 1-10 had significantly higher median Δdist-AICc values in LIZARDS (**table 2**).

**Figure 2:**
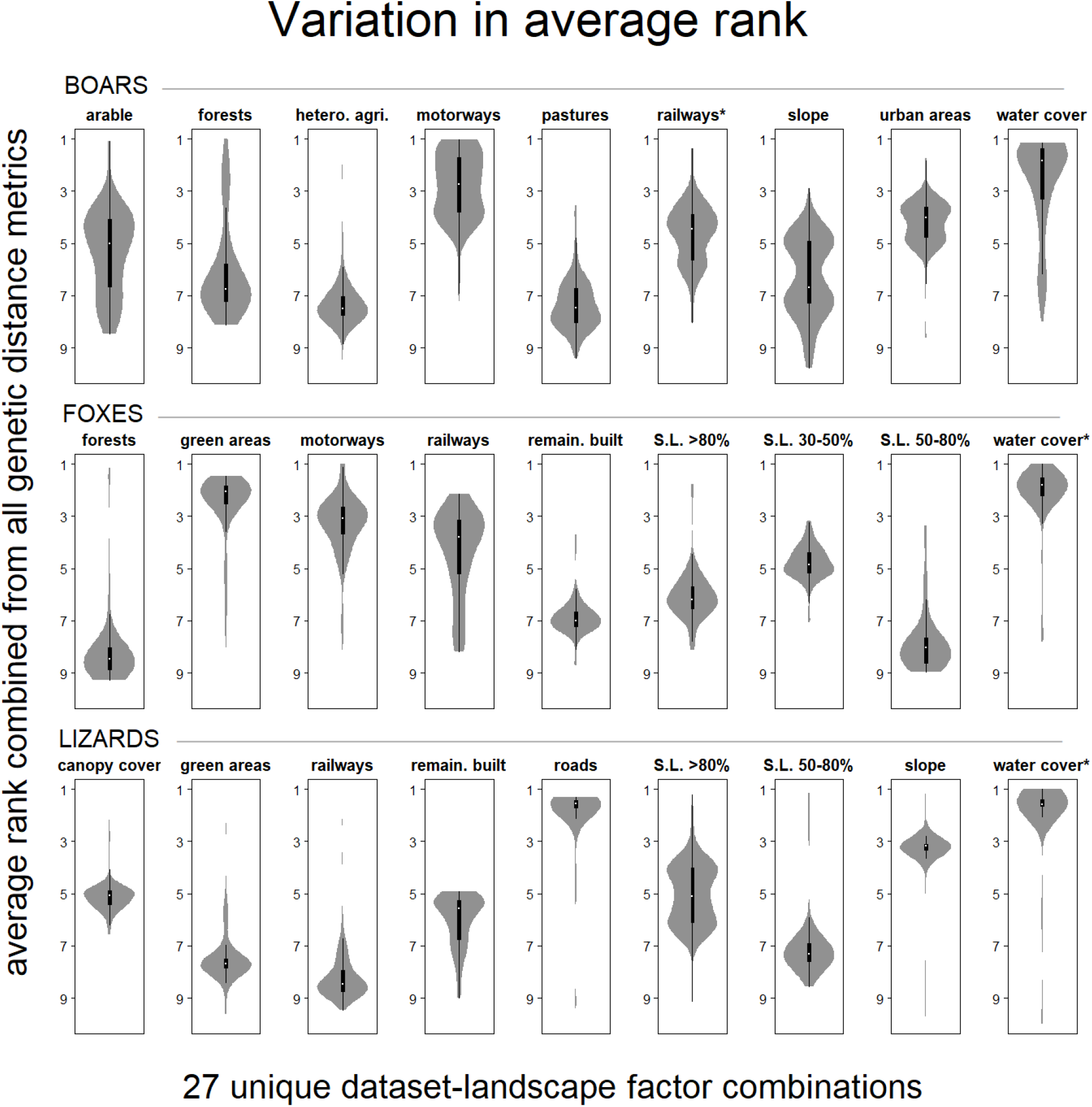
Average rank of models of all landscape factors after b ootstrapping. Highest ranking models based on Δdist-AICc values within datasets are marked with an asterisk.

**Figure 3:**
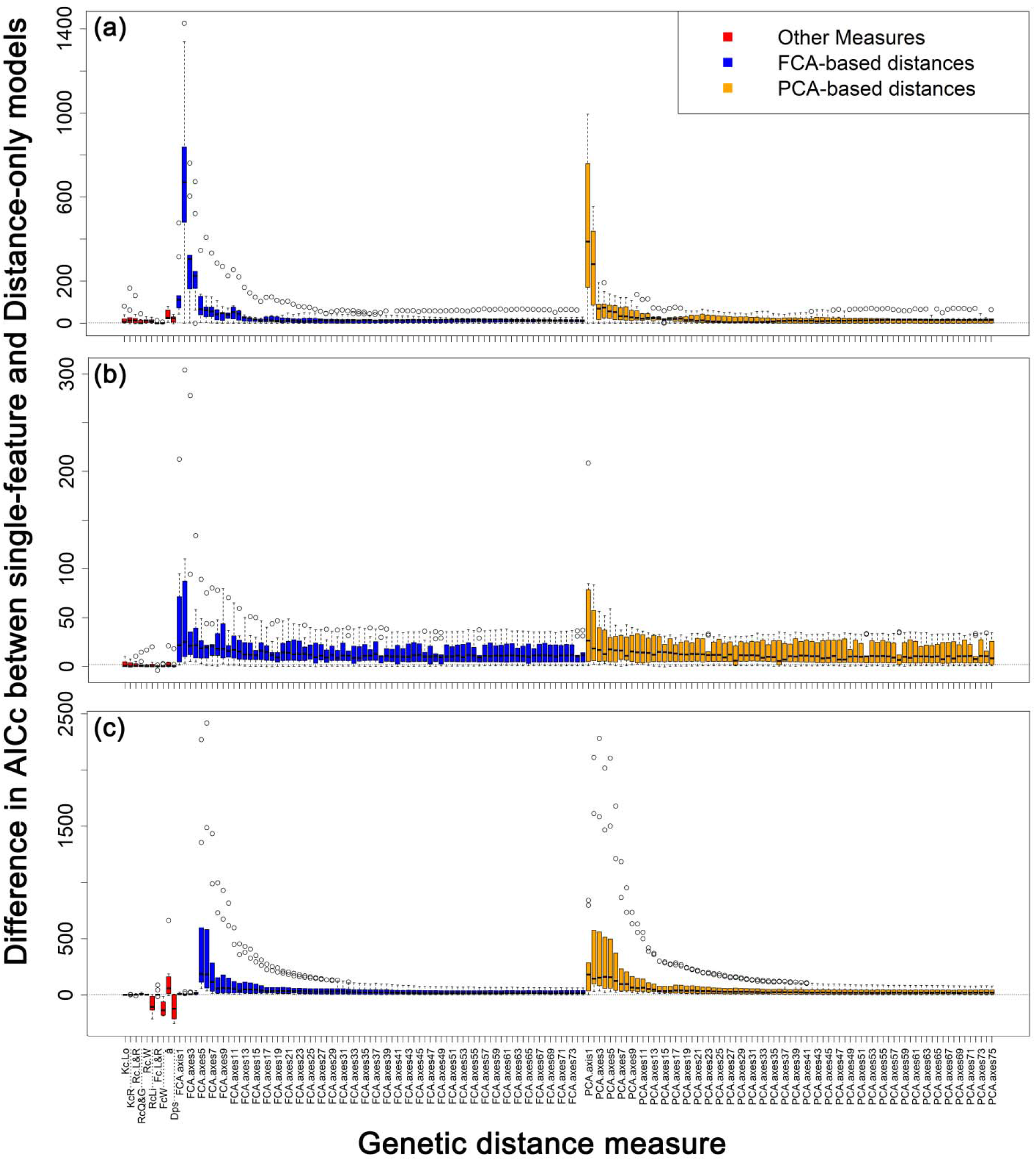
Δdist-AICc values of all 8640 optimizations for BOARS (a), FOXES (b) and LIZARDS (c).

**Table 2:** Comparison of Δdist-AICc values of optimized models using genetic distance metrics derived from FCA and PCA axes 1-10, FCA and PCA axes 11-75 and Other Metrics. Median distAICc-values of models (A) and results of tests of FCA and PCA axes 1-10 against FCA and PCA axes 11-75, Other Metrics and against each other using Wilcoxon rank sum test (B).

| *Genetic distance metrics* | *BOARS* | *FOXES* | *LIZARDS* |
| --- | --- | --- | --- |
| **FCA 1-10** | 72.74 | 20.41 | 27.46 |
| **PCA 1-10** | 61.30 | 16.44 | 129.67 |
| **FCA 11-75** | 14.08 | 11.06 | 19.22 |
| **PCA 11-75** | 14.96 | 10.40 | 21.56 |
| **Other Metrics** | 5.42 | 0.31 | -0.05 |

(B)
| *comparison* | *BOARS* | *FOXES* | *LIZARDS* |
| --- | --- | --- | --- |
| **FCA_1-10 vs FCA_11_75** | W=43919, $P<0.001$ | W=34396, $P<0.001$ | W=30045, $P=0.031$ |
| **PCA_1-10 vs PCA_11_75** | W=39657, $P<0.001$ | W=32045, $P<0.001$ | W=42938, $P<0.001$ |
| **FCA_1-10 vs Other Metrics** | W=7101, $P<0.001$ | W=7645, $P<0.001$ | W=7086, $P<0.001$ |
| **PCA_1-10 vs Other Metrics** | W=6589, $P<0.001$ | W=7430, $P<0.001$ | W=7739, $P<0.001$ |
| **FCA_1-10 vs PCA_1-10** | W=4789, $P=0.035$ | W=4690, $P=0.067$ | W=2431, $P<0.001$ |

No single genetic distance metric consistently yielded models with the highest Δdist-AICc values, within and across datasets (**figure 3**; **table 1 & S2**). The highest-ranking model within each dataset was based on an FCA derived genetic distance metric (based on two axes in BOARS and FOXES and on six axes in LIZARDS; **table 1**). However, when considering both the first ten axes of FCA- or PCA-based metrics, visual inspection of median and maximum Δdist-AICc values (**figures 3 & S10-S12**) allowed the identification of a single metric at which the highest or close-to-the-highest Δdist-AICc values were obtained for at least seven of the nine landscape features in each dataset (**figures S10-S12)**.

Marginal R^2^ values (mR^2^) of models based on FCA axes 1-10 were significantly lower than those and based on PCA axes 1-10 in all datasets (BOARS: Wilcoxon rank sum test: W=3119, *P*=0.0078; FOXES: W=2962, *P*=0.0019; LIZARDS: W=1599, *P*<0.0001). At the same time, mR^2^ values of the first ten axes of both FCA and PCA were a multiple of the values obtained by models using the ten Other Metrics or axes 11-75 of FCA and PCA, frequently up to an order of magnitude greater (**table S3**).

### Influence of genetic distance metrics on results of landscape factors and optimized resistances

The resistances of optimized models of each landscape factor across all 160 genetic distance metrics varied substantially (**figures S13-15**; **table S3 & S4**). With a focus on the 25 binary landscape factor models used in analyses (which excludes the continuous model of slope in BOARS and LIZARDS) models of only two landscape factors were consistently optimized with resistances that had the same directionality of effecting gene flow, irrespective of genetic distance metric. In the remaining 23 landscape factors, resistances of optimized models switched between facilitating or impeding gene flow, depending on the genetic distance metric utilized in the analysis (**table S4**). Within the genetic distance metrics derived from the first ten axes of both FCA and PCA, such switches happened less frequently (FCA: three cases; PCA: no cases) than in the ten Other Metrics or genetic distance metrics derived from axes 11-75 of FCA and PCA (**table S4**).

## Discussion

Landscape genetic approaches hold great promise for conservation applications, as they can provide scientific guidance for conservation management in general and present an empirical basis for the optimal design of movement corridors in specific (Keller et al., 2014). However, to achieve these goals, landscape genetic approaches need to deliver ecologically meaningful and methodologically robust results. A large body of simulation studies already shows that conclusions of landscape genetic studies can be impacted by a large variety of methodological and statistical choices (Peterman et al., 2019; Peterman, Connette, Semlitsch, & Eggert, 2014; Richardson, Brady, Wang, & Spear, 2016; Shirk, Landguth, & Cushman, 2017a & 2017b). Surprisingly, the effect of genetic distance metrics on landscape genetic inferences has received relatively little attentions (but see Shirk, Landguth, & Cushman, 2017a; Kimmig et al. 2020). Based on a thorough analysis of three individual-based, empirical datasets with the R-package ResistanceGA, we here provide clear evidence for a strong impact of the choice of genetic distance metric on landscape genetic inferences (**figures 2 & 3**).

Our results strongly suggest that there are clear differences in AICc-based model support, marginal R^2^ (mR^2^)-based model fit and directionality of inferred landscape resistances between different genetic distance metrics. Genetic distance metrics based on the first ten axes of a FCA or PCA outperform the more commonly used Other Metrics, including the widely used proportion of shared alleles (D_PS_) and other kinship- or relatedness-coefficients (**table 2**). Highest maximum Δdist-AICc values (for all but one landscape factor), highest median Δdist-AICc values, highest maximum and median mR^2^ values as well as fewest switches in directionality of resistances were all obtained when utilizing FCA and PCA-based metrics that calculated genetic distances based on between 1-10 axes. In many cases, the best-supported models obtained from the first ten axes of FCA or PCA had Δdist-AICc and mR^2^ values an order of magnitude greater than of models utilizing any of the Other Metrics or axes 11-75 of FCA and PCA (**figure 3**; **table S3**).

It is perhaps not surprising that FCA/PCA-based measures perform well, as both methods emphasise differences better than metrics that weight all loci equally (such as all Other Metrics). Unsurprisingly, other studies have also foud FCA/PCA-based measures to provide better model support or fit than other genetic distance metrics, based on Mantel’s test correlation (Shirk, Wallin, Cushman, Rice, & Warheit, 2010), AICc values (Shirk et al. 2017a) or r^2^ (Mignotte et al., 2020). In the majority of cases, higher maximum Δdist-AICc values were obtained for each landscape factor with FCA- rather than PCA-based metrics. FCA, which assumes data to consist of multistate categorical variables, ought to be a priori more suitable for the analysis of genetic data than PCA, which assumes continuous, normally distributed data. At the same time, model fit evaluated by mR^2^ values of models was significantly higher in models utilizing the first ten axes of PCA as opposed to those of FCA, an observation that is not readily explainable and requires further study.

While we are focussing on single-feature gene flow models in the present paper, a comprehensive understanding of landscape connectivity from the species’ perspective, i.e. functional connectivity, requires simultaneous consideration of different landscape features (Peterman & Pope 2020). When using ResistanceGA, the optimal approach for performing a multivariate analysis can be far from trivial – especially when overlapping linear landscape variables are considered (Kimmig et al. 2020) – and the role of genetic distance metrics on multivariate analyses, more generally, needs further investigation, which is beyond the scope of this study. Nevertheless, also multivariate analyses require a single genetic distance metric to evaluate each composite model under consideration. This study shows that there is no single genetic distance metric that consistently yielded the highest model support and fit across datasets and single-feature landscape models within datasets. In the case of the FCA/PCA-based metrics, however, visual inspection of maximum and median Δdist-AICc values suggested that it is possible to identify a metric based on a specific number of axes that gives rise to models with the highest or close-to-the-highest Δdist-AICc values for at least seven of the nine individual landscape features tested in each dataset. We thus recommend that authors focus on the first ten FCA and/or PCA axes to identify an ‘optimal’ number of axes that maximises model support for all, or most, landscape features and which can be used as the genetic distance metric for all analyses.

For the empirical datasets presented here we have no knowledge regarding the true extent to which landscape factors are influencing gene flow. However, all drawn conclusion rest on AICc-based model support and mR -based model fit. Optimisation of resistance models followed by AICc ranking of LME models is methodologically sound and especially suited for pairwise distance matrices (Shirk et al., 2017b). However, simulations by Winiarski et al. (2020) have suggested an elevated type I error when applying the AICc-model selection approach implemented in ResistanceGA to individual-based, spatially correlated alternative landscape resistance surfaces. Results by Kimmig et al. (2020) suggest a stepwise multivariate analysis might identify composite models which exclude features that do not truly influence gene flow. In this approach, single-feature landscape models are sequentially added to a composite model and retained only if their addition improved model support (ΔAICc > 2). Therefore, while using FCA/PCA based genetic distance metrics may lead to an increased type I error in univariate analyses, a subsequent multivariate analysis can, in all likelihood, identify a composite model, which excludes such landscape models.

Recent landscape genetic simulations have adopted a genetic distance metric derived from 64 PCA axes (Winiarski et al. (2020) or Pope & Peterman (2020)), due to simulation results by Shirk et al. (2017a). These showed that a metric derived from 64 PCA axe allowed to identify the true landscape resistance surface most frequently, under various simulation scenarios. Our results, however, demonstrate that when using genetic distance metrics based on a number of FCA/PCA axes above ten, Δdist-AICc and mR^2^ values decrease and variation in the directionality of optimised resistances increases. The use of Other Metrics came to similar results relative to models based on the first ten FCA/PCA axes. This corroborates that some genetic distance metrics inferred erroneous resistance values and we therefore call for a more cautious interpretation of resistance values, generally, but especially when results are not based on genetic distance metrics derived from the first ten axes of FCA/PCA (e.g. Beninde et al. 2018) and they appear counter-intuitive in light of the ecology of species, (e.g. Peterman et al., 2014). The resistance values inferred for each landscape feature in the present study appear a priori to be meaningful (table S2; for FOXES see also Kimmig et al. 2020).

One explanation for the very different conclusions drawn here and the results by Shirk et al. (2017a) may reside in the parameter settings used for some of their simulations. Their results showed that many other genetic distance metrics, including D_PS_ and 4 or 16 PCA axes, had similar model selection accuracy as 64 PCA axes when simulations allowed individuals to disperse up to 20% of the largest pairwise Euclidean distance between individuals only or sample size was high (N>1000). However, when individuals were allowed to disperse up to 100% of the largest pairwise Euclidean distance between individuals and sample size was low (N=200), relative model selection accuracy of all other genetic distance metrics decreased and was outperformed by that derived from 64 PCA axes. We presume that authors have therefore adopted the genetic distance metric derived form 64 PCA axes, as a safe choice to maximize model selection accuracy across various scenarios. However, arguably, the extent of landscape genetic studies is frequently a multiple of the maximum dispersal distance of target species and, therefore, the conclusion that 64 PCA axes deliver the most suitable genetic distance metric may be misleading.

Many open questions remain regarding the performance of genetic distance metrics in inferring landscape genetic resistance. Simulations have suggested an increase of model selection accuracy with lower variance in genetic distances (Winiarski et al. 2020). However, the sample variance of the principal components of eigenvector-based multivariate analysis increases with decreasing sample size (Trochimczyk & Chayes 1977, Björklund 2019). The applicability of conclusions presented needs further study for datasets with small numbers of samples or loci. Perhaps the preliminary study called for here should explore more than the first ten axes in studies with such datasets. Furthermore, the conclusions presented here are based on microsatellite loci. Principally, single-nucleotide polymorphisms (SNPs), especially when used in their thousands, ought to allow more precise estimation of genetic relatedness, fraternity and dissimilarity (e.g. Lemopoulos et al 2019, Galla et al. 2020). It thus remains to be seen whether the results presented here will be confirmed for studies using higher-resolution genomic makers, or whether, because of reduced sampling variance, the choice of genetic distance metric matters less when working with SNPs. Finally, we largely focus on categorical variables (25/27 landscape factors) and further research is required to assess the accuracy of conclusions also for continuous surfaces.

### Conclusions

We believe that our findings are of wide-ranging significance, also beyond landscape genetic analyses, for any study utilizing genetic distances of individuals for inferences of, e.g. fragmentation, connectivity, corridors, or, more generally, ecological and evolutionary patterns. While empirical datasets have no knowledge of true, underlying driving factors, simulation studies are limited in the number conditions and the parameter space it can explore, which can be especially disadvantageous when not matched to the experimental design of empirical studies. While it is important to corroborate the conclusions drawn here using simulated datasets, with known underlying resistances values, it is necessary to point out that, likewise, results from simulation studies should be subjected to the same scrutiny and corroboration by empirical datasets.

Many questions remain unanswered and need to be explored further to increase robustness and reliability of landscape genetic inferences, in particular. However, the results presented here emphasize an imperative preliminary step in analyses, testing the effect of genetic distance metrics on model performance when utilizing genetic distances of individuals. Based on the assembled evidence, we recommend exploring the first ten axes of FCA and/or PCA to identify a genetic distance metric that gives rise to models with the highest or close-to-the-highest Δdist-AICc values most frequently. It remains a challenge to infer landscape genetic resistances of empirical datasets, and we here present an approach to handle uncertainty in such inferences in the most robust way possible. Our results indicate that, if this preliminary step is not conducted, the importance of landscape factors on gene flow may remain concealed by the arbitrary choice of genetic distance metrics, preventing an accurate interpretation of the underlying ecological, driving pattern.

## Supporting information

All supplementary materials

## Acknowledgements

We would like to thank Alain Licoppe, Sabine Bertouille and the local Services of the Nature and Forest Department (General Directorate for Agriculture, Natural Resources and Environment of the Public Service of Wallonia) for sample collection and MC Flamand (Louvain Institute of Biomolecular Science and Technology, Université catholique de Louvain) for genotyping of the wild boar samples. We would like to thank Paul Wilmes, Cédric Laczny and Maharshi Vyas for granting us access to the computing cluster of the Systems Ecology research group at the University of Luxembourg. JB work was funded by a DFG grant (BE 6887/1-1). The work was support by an internal grant from the Musée National d’Histoire Naturelle, Luxembourg.

## Notes

### Competing Interest Statement

The authors have declared no competing interest.

