## Supplementary material for "Using empirical datasets to quantify uncertainty in inferences of landscape genetic resistance due to variation of individual-based genetic distance metrics": All supplementary materials

Contents:

**Table S1:** Characteristics of the spatial datasets used in this study.
**Table S2:** Results for all 27 dataset-landscape factor combinations models per dataset
**Table S3:** Range of marginal $R^2$-values
**Table S4:** Directionality of resistance score in optimized models of landscape factors
**Figure S1:** Landscape models of BOARS.
**Figure S2:** Landscape models of FOXES.
**Figure S3:** Landscape models of LIZARDS.
**Figure S4:** Histogram of the ΔAICc differences between two runs of optimization
**Figure S5:** BOARS results of both runs for FCA/PCA axes 1-75
**Figure S6:** FOXES results of both runs for FCA/PCA axes 1-75
**Figure S7:** LIZARDS results of both runs for FCA/PCA axes 1-75
**Figure S8:** Differences in Δdist-AICc between initial and bootstrapping results
**Figure S9:** Difference in Δdist-AICc of models with resistances >500.
**Figure S10:** BOARS Δdist-AICc of Other Metrics and FCA/PCA axes 1-10.
**Figure S11:** FOXES Δdist-AICc of Other Metrics and FCA/PCA axes 1-10.
**Figure S12:** LIZARDS Δdist-AICc of Other Metrics and FCA/PCA axes 1-10.
**Figure S13:** Resistances Other Metrics and FCA/PCA axes 1-10 for BOARS.
**Figure S14:** Resistances Other Metrics and FCA/PCA axes 1-10 for FOXES
**Figure S15:** Resistances Other Metrics and FCA/PCA axes 1-10 for LIZARDS

**Table S1 Characteristics of the spatial datasets used in this study.**

| DATASET | LANDSCAPE FEATURE | SOURCE | CLASSES | SOURCE |
|---|---|---|---|---|
| **BOARS** | *arable* | Corine Land Cover 2018 | 2.1 Arable land; 2.2 Permanent crops | 1 |
| **BOARS** | *forests* | Corine Land Cover 2018 | 3. Forest and semi-natural areas | 1 |
| **BOARS** | *hetero. agri.* | Corine Land Cover 2018 | 2.4 Heterogeneous agricultural areas | 1 |
| **BOARS** | *motorways* | Open Street Map | Roads | 2 |
| **BOARS** | *pastures* | Corine Land Cover 2018 | 2.3 Pastures | 1 |
| **BOARS** | *railways* | Open Street Map | Railways | 2 |
| **BOARS** | *urban areas* | Corine Land Cover 2018 | 1. Artificial Surfaces | 1 |
| **BOARS** | *slope* | EU-DEM v1.1 | EU Digital Elevation model | 3 |
| **BOARS** | *water cover* | Corine Land Cover 2018 | OSM: Waterways; CLC: 5. Water bodies | 1, 2 |
| **FOXES** | *forests* | ATKIS | Waldflaechen | 4 |
| **FOXES** | *green areas* | ATKIS | Ackerland, Brachen, Gartenland, Grünland | 4 |
| **FOXES** | *motorways* | ATKIS & OSM | ATKIS: Strassen, OSM: Roads | 2, 4 |
| **FOXES** | *railways* | ATKIS | Flächenhafte Verkehrsobjekte | 4 |
| **FOXES** | *remain. built-up* | Urban Atlas 2012 | 1. Artificial surfaces (except classes 11100, 11210 & 11220; see below) | 5 |
| **FOXES** | *S.L. 30-50%* | Urban Atlas 2012 | 11220 Discontinuous medium density urban fabric (S.L. 30%-50%) | 5 |
| **FOXES** | *S.L. 50-80%* | Urban Atlas 2012 | 11210 Discontinuous dense urban fabric (S.L. 50%-80%) | 5 |
| **FOXES** | *S.L. >80%* | Urban Atlas 2012 | 11100 Continuous urban fabric (S.L.: >80%) | 5 |
| **FOXES** | *water cover* | ATKIS | Flächenhafte Gewässer | 4 |
| **LIZARDS** | *canopy cover* | Urban Atlas 2012 & Street Tree Layer 2018 | Urban Atlas: 3. Forest (natural and plantation); Street Tree Layer 2018 | 5, 6 |
| **LIZARDS** | *green areas* | Urban Atlas 2012 | 2. Agricultural and Semi-natural | 5 |
| **LIZARDS** | *railways* | OSM | Railways | 2 |
| **LIZARDS** | *roads* | OSM | Roads | 2 |
| **LIZARDS** | *remain. built-up* | Urban Atlas 2012 | 1. Artificial surfaces (except classes 11100 & 11210; see below) | 5 |
| **LIZARDS** | *S.L. 50-80%* | Urban Atlas 2012 | 11210 Discontinuous dense urban fabric (S.L. 50%-80%) | 5 |
| **LIZARDS** | *S.L. >80%* | Urban Atlas 2012 | 11100 Continuous urban fabric (S.L.: >80%) | 5 |
| **LIZARDS** | *slope* | DTM Germany | Digital LiDAR-Terrain Model of Germany: Rhineland-Palatinate, 20 meters | 7 |
| **LIZARDS** | *water cover* | OSM | Waterways | 2 |

https://land.copernicus.eu/pan-european/corine-land-cover/clc2018; 2: https://download.geofabrik.de; 3; https://land.copernicus.eu/imagery-in-situ/eu-dem/eu-dem-v1.1; 4: Gruenreich, D. (1992). ATKIS-a topographic information system as a basis for GIS and digital cartography in Germany. From Digital Map Series to Geo-Information Systems, Geologisches Jahrbuch Series A. Hannover, Germany: Federal Institute of Geosciences and Resources; 5: https://land.copernicus.eu/local/urban-atlas/urban-atlas-2012; 6: https://land.copernicus.eu/local/urban-atlas/street-tree-layer-stl-2018; 7: https://bit.ly/dtm-germany-rheinland-pfalz-20m (all links accessed 23/12/2020).

**Table S2: Results for all 27 dataset-landscape factor combinations models per dataset**

| ***dataset*** | *Landscape factor* | *Genetic distance metric* | *Highest-model Δdist-AICc* | *Highest-model resistance* | *Lowest-model Δdist-AICc* | *Overall minimum resistance* | *Overall maximum resistance* |
|---|---|---|---|---|---|---|---|
| **BOARS** | railways | FCA 2 axes | 1427.20 | 61.61 | 1.08 | -23.16 | 159.04 |
| **BOARS** | motorways | FCA 2 axes | 1339.82 | 59.77 | 0.16 | -66.29 | 59.77 |
| **BOARS** | forests | PCA 1 axes | 994.63 | -38.08 | -0.03 | -93.30 | 2.14 |
| **BOARS** | arable | FCA 2 axes | 748.79 | 9.60 | -0.06 | -4.20 | 20.36 |
| **BOARS** | slope | FCA 2 axes | 670.19 | NA | -1.17 | NA | NA |
| **BOARS** | pastures | FCA 2 axes | 517.01 | -6049.46 | -0.05 | -6049.46 | 52.02 |
| **BOARS** | urban areas | FCA 2 axes | 480.57 | 18.92 | 0.25 | -1.07 | 34.56 |
| **BOARS** | water cover | PCA 1 axes | 171.33 | -22.74 | 0.19 | -177.93 | 1.17 |
| **BOARS** | hetero. agri. | FCA 4 axes | 63.99 | 3.82 | -0.32 | -5.70 | 499.94 |
| **FOXES** | water cover | FCA 2 axes | 304.09 | 378.06 | -0.52 | -81.96 | 378.06 |
| **FOXES** | railways | FCA 2 axes | 110.54 | -303.29 | -0.49 | -3352.76 | 12.03 |

| | | | | | | | |
|---|---|---|---|---|---|---|---|
| FOXES | green areas | FCA 3 axes | 94.70 | -9993.09 | -0.35 | -9998.26 | 3.58 |
| FOXES | motorways | PCA 1 axes | 55.89 | -303.29 | -0.09 | -7580.02 | 1.14 |
| FOXES | S.L. 50-80% | FCA 3 axes | 30.29 | -3019.94 | -0.36 | -4872.75 | 1.93 |
| FOXES | S.L. 30-50% | PCA 1 axes | 26.72 | -5876.63 | -0.15 | -9932.10 | -1.01 |
| FOXES | S.L. >80% | FCA 8 axes | 13.80 | 9.26 | -0.15 | -566.81 | 11.72 |
| FOXES | forests | Lynch | 10.63 | -1.92 | -0.10 | -1.92 | 8.54 |
| FOXES | remain. built-up | FCA 10 axes | 9.72 | 9.55 | -4.00 | -2.03 | 499.97 |
| LIZARDS | water cover | FCA 6 axes | 2416.41 | 499.95 | -254.96 | -29.60 | 499.97 |
| LIZARDS | roads | PCA 2 axes | 1611.40 | 500.00 | -255.31 | -1.19 | 500.00 |
| LIZARDS | slope | FCA 5 axes | 596.68 | NA | -213.71 | NA | NA |
| LIZARDS | S.L. >80% | PCA 4 axes | 255.09 | 500.00 | -15.37 | -8.36 | 500.00 |
| LIZARDS | green areas | PCA 1 axes | 186.86 | -67.20 | -186.92 | -256.76 | 1.05 |
| LIZARDS | canopy cover | FCA 5 axes | 183.99 | -94.25 | -149.20 | -94.25 | -1.10 |

| | | | | | | | |
|---|---|---|---|---|---|---|---|
| **LIZARDS** | railways | FCA 5 axes | 111.97 | -65.54 | -106.91 | -65.54 | 2.06 |
| **LIZARDS** | S.L. 50-80% | PCA 2 axes | 96.92 | 21.00 | -46.95 | -1.06 | 172.46 |
| **LIZARDS** | remain. built-up | FCA 5 axes | 56.59 | 7.64 | -144.36 | -1.27 | 7.64 |

**Table S3: Range of marginal $R^2$-values of models using genetic distance metrics derived from the first 10 axes and higher of FCA and PCA axes and of Other Metrics (median in parentheses; tested using Wilcoxon rank sum test).**

| ***Genetic distance metric*** | *BOARS* | *FOXES* | *LIZARDS* |
|---|---|---|---|
| **FCA_1-10** | 0.0054-0.3329 (0.028) | 0.0101-0.58 (0.0343) | 0.0005-0.5007 (0.0117) |
| **PCA_1-10** | 0.0146-0.3838 (0.0381) | 0.0181-0.603 (0.0438) | 0.0116-0.6465 (0.047) |
| **FCA_11-75** | 0.0019-0.0418 (0.0044) | 0.0038-0.2181 (0.0155) | 0.0023-0.3724 (0.0087) |
| **PCA_11-75** | 0.0024-0.0372 (0.0063) | 0.0059-0.2242 (0.0235) | 0.0032-0.4446 (0.0119) |
| **Other Metrics** | <0.001-0.0382 (0.0182) | 0.0008-0.0531 (0.014) | 0.001-0.2228 (0.027) |

**Table S4: Directionality of resistance score in optimized models of landscape factors, separately for the first 10 axes of FCA and PCA axes, higher axes of FCA and PCA and Other Metrics (barrier=barrier to gene flow; conduit: facilitator of gene flow; switched=both barrier and conduit; for slope the number of different transformations is given).**

| *Landscape factor* | *FCA 1-10* | *PCA 1-10* | *FCA 11-75* | *PCA 11-75* | *Other Metrics* |
|---|---|---|---|---|---|
| **arable (Boars)** | barrier | barrier | switched | switched | switched |
| **forests (Boars)** | conduit | conduit | switched | switched | switched |
| **hetero. agri. (Boars)** | barrier | barrier | switched | switched | switched |
| **motorways (Boars)** | barrier | barrier | barrier | barrier | switched |
| **pastures (Boars)** | switched | conduit | switched | conduit | switched |
| **railways (Boars)** | switched | barrier | barrier | barrier | switched |
| **slope (Boars)** | 2 | 1 | 2 | 1 | 3 |
| **urban areas (Boars)** | barrier | barrier | barrier | barrier | switched |
| **water cover (Boars)** | conduit | conduit | conduit | conduit | switched |
| **forests (Foxes)** | barrier | barrier | switched | barrier | switched |
| **green areas (Foxes)** | conduit | conduit | conduit | conduit | switched |
| **motorways (Foxes)** | conduit | conduit | conduit | conduit | switched |
| **railways (Foxes)** | conduit | conduit | switched | switched | switched |
| **remain. built-up (Foxes)** | barrier | barrier | barrier | barrier | switched |
| **S.L. >80% (Foxes)** | barrier | barrier | barrier | barrier | switched |
| **S.L. 30-50% (Foxes)** | conduit | conduit | conduit | conduit | conduit |
| **S.L. 50-80% (Foxes)** | conduit | conduit | conduit | conduit | switched |
| **water cover (Foxes)** | barrier | barrier | barrier | barrier | switched |

| | | | | | |
|---|---|---|---|---|---|
| **canopy cover (Lizards)** | conduit | conduit | conduit | conduit | conduit |
| **green areas (Lizards)** | conduit | conduit | conduit | conduit | switched |
| **railways (Lizards)** | conduit | conduit | conduit | conduit | switched |
| **remain. built-up (Lizards)** | barrier | barrier | barrier | barrier | switched |
| **roads (Lizards)** | barrier | barrier | barrier | barrier | switched |
| **S.L. >80% (Lizards)** | barrier | barrier | barrier | barrier | switched |
| **S.L. 50-80% (Lizards)** | barrier | barrier | barrier | barrier | switched |
| **slope (Lizards)** | 3 | 1 | 1 | 1 | 3 |
| **water cover (Lizards)** | switched | barrier | barrier | barrier | switched |

**Figures S1:** Landscape models of BOARS.

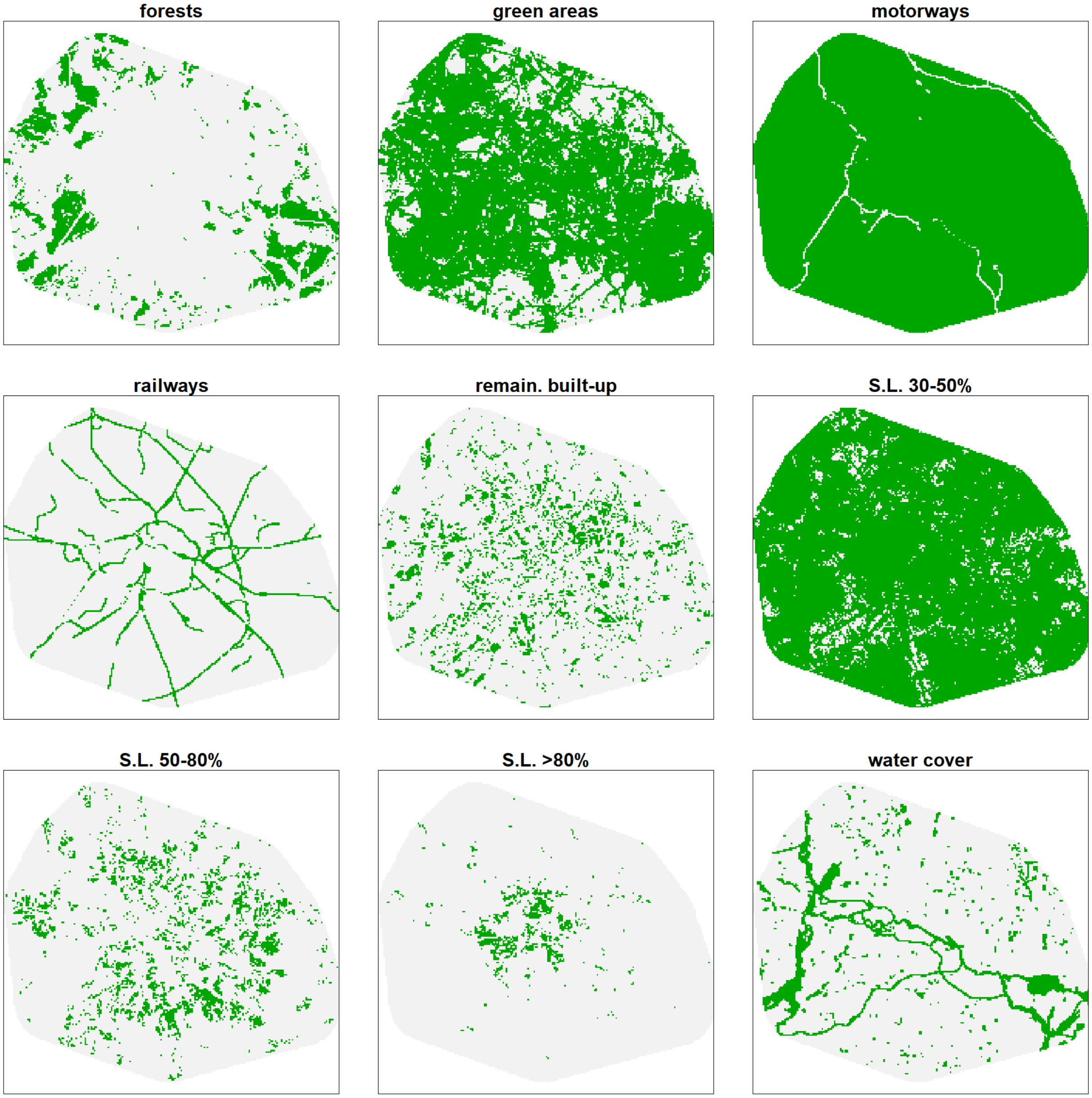

**Figures S2:** Landscape models of FOXES.

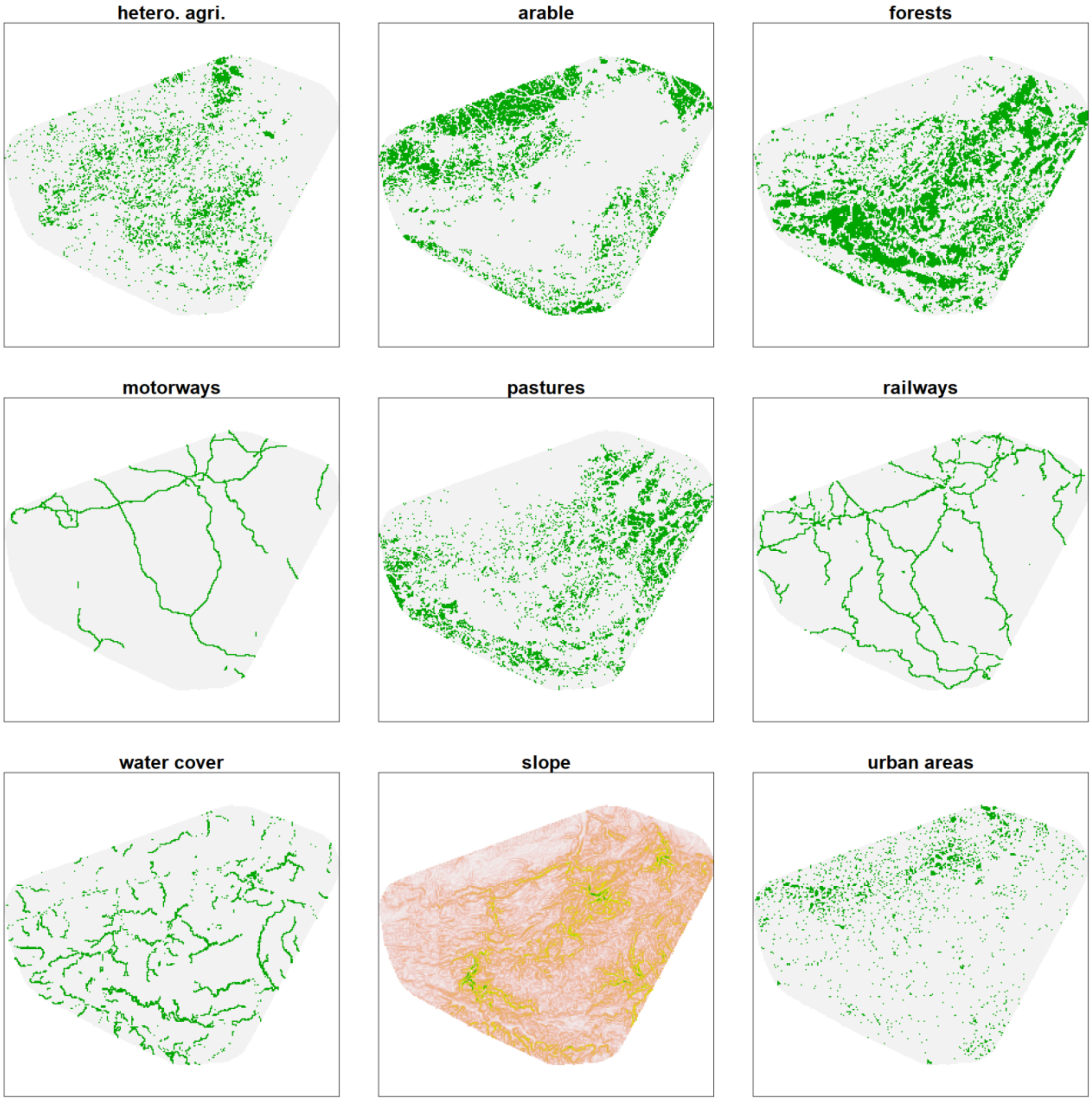

**Figures S3:** Landscape models of LIZARDS.

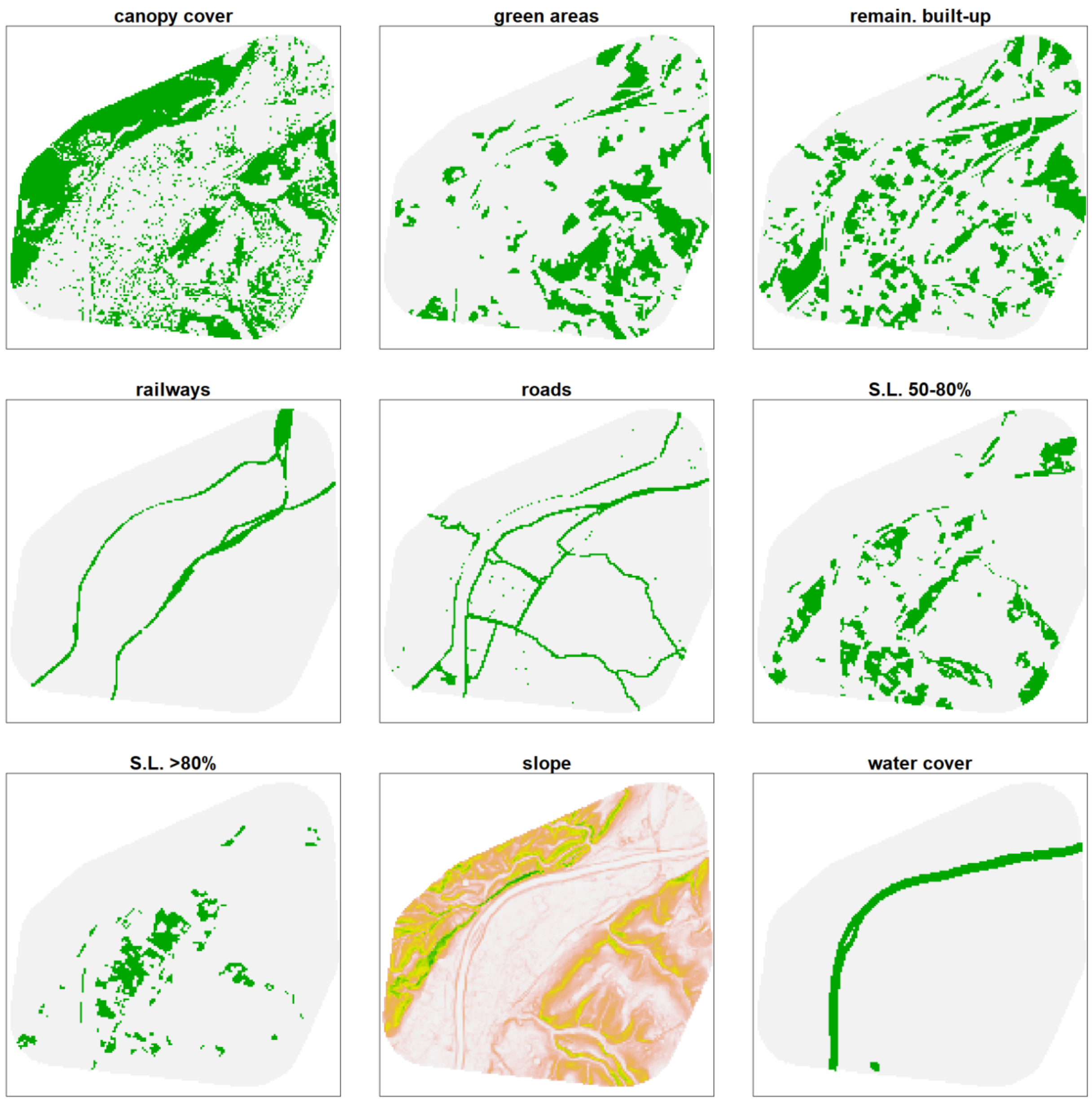

**Figure S4:** Histogram of the ΔAICc differences between the models of two runs of optimization for identical dataset-landscape factor-genetic distance metric combinations.

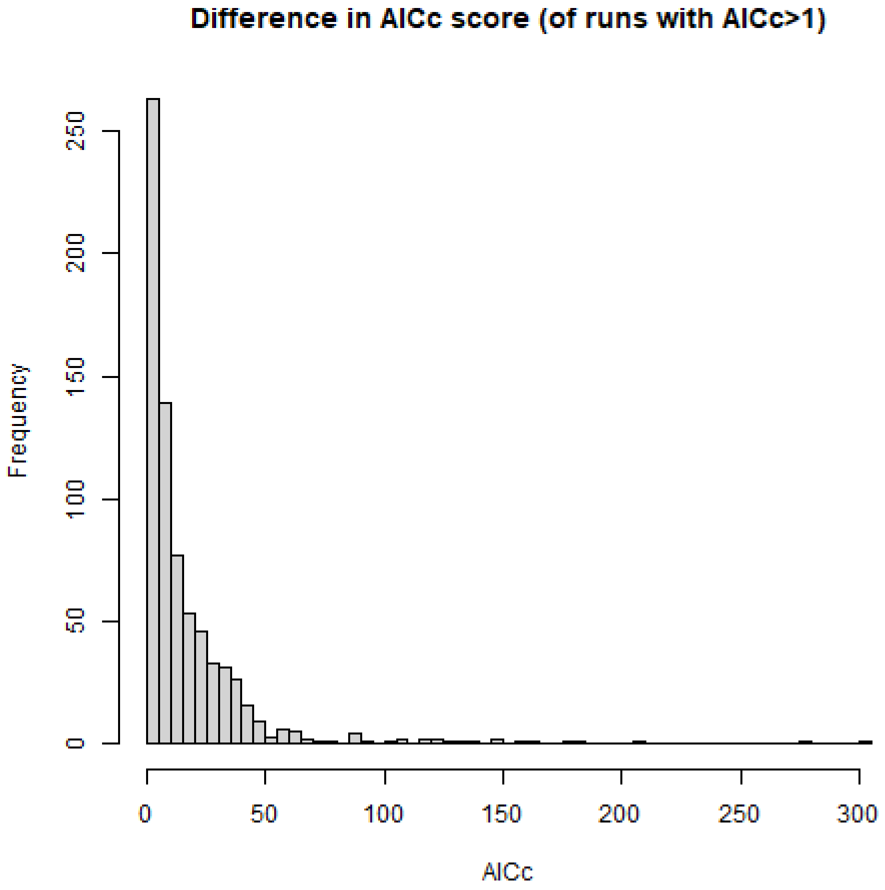

**Figures S5:** BOARS results of both runs of each landscape factor for all genetic distances metrics derived from FCA/PCA axes 1-75.

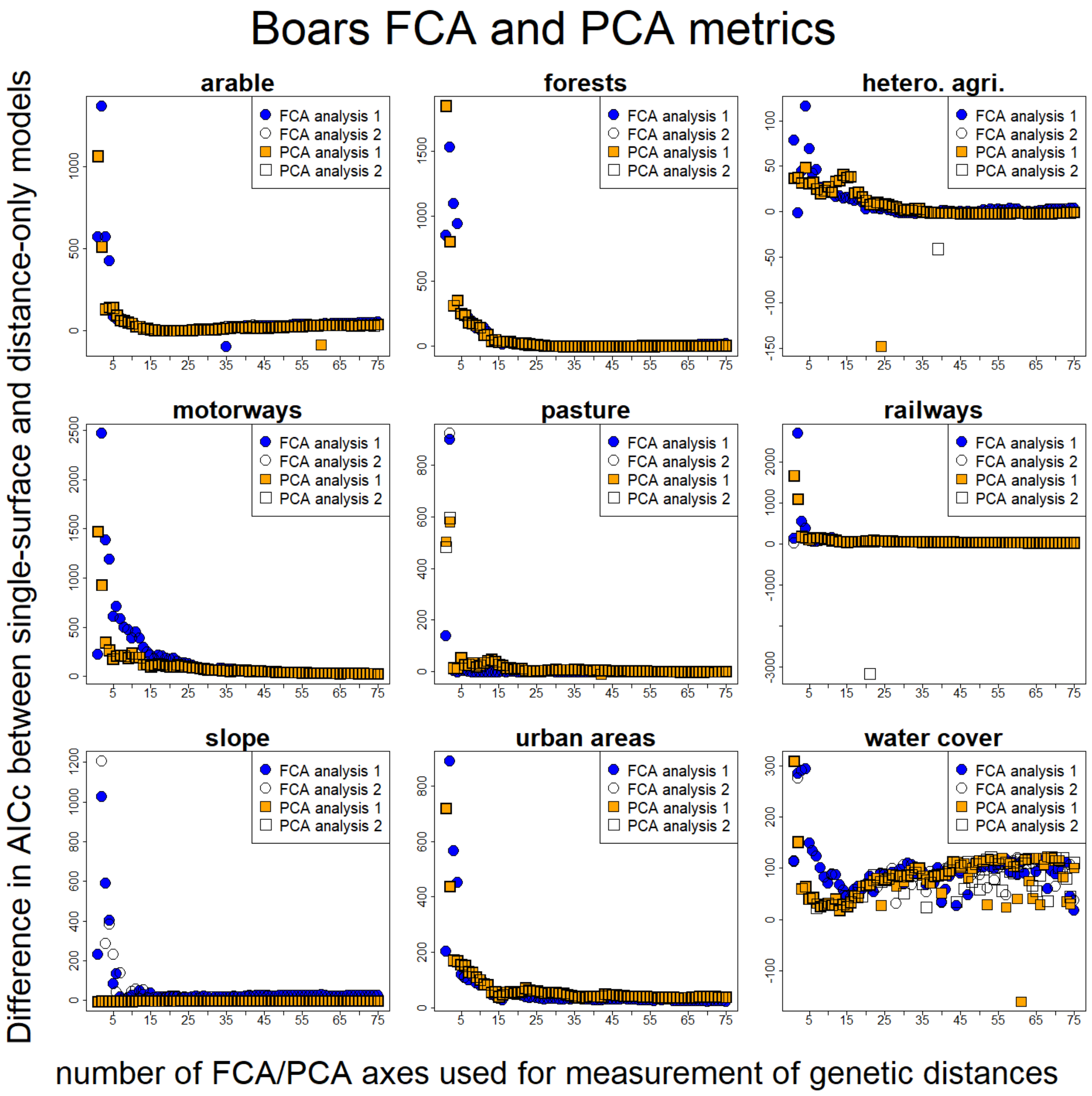

**Figures S6: FOXES results of both runs of each landscape factor for all genetic distances metrics derived from FCA/PCA axes 1-75**

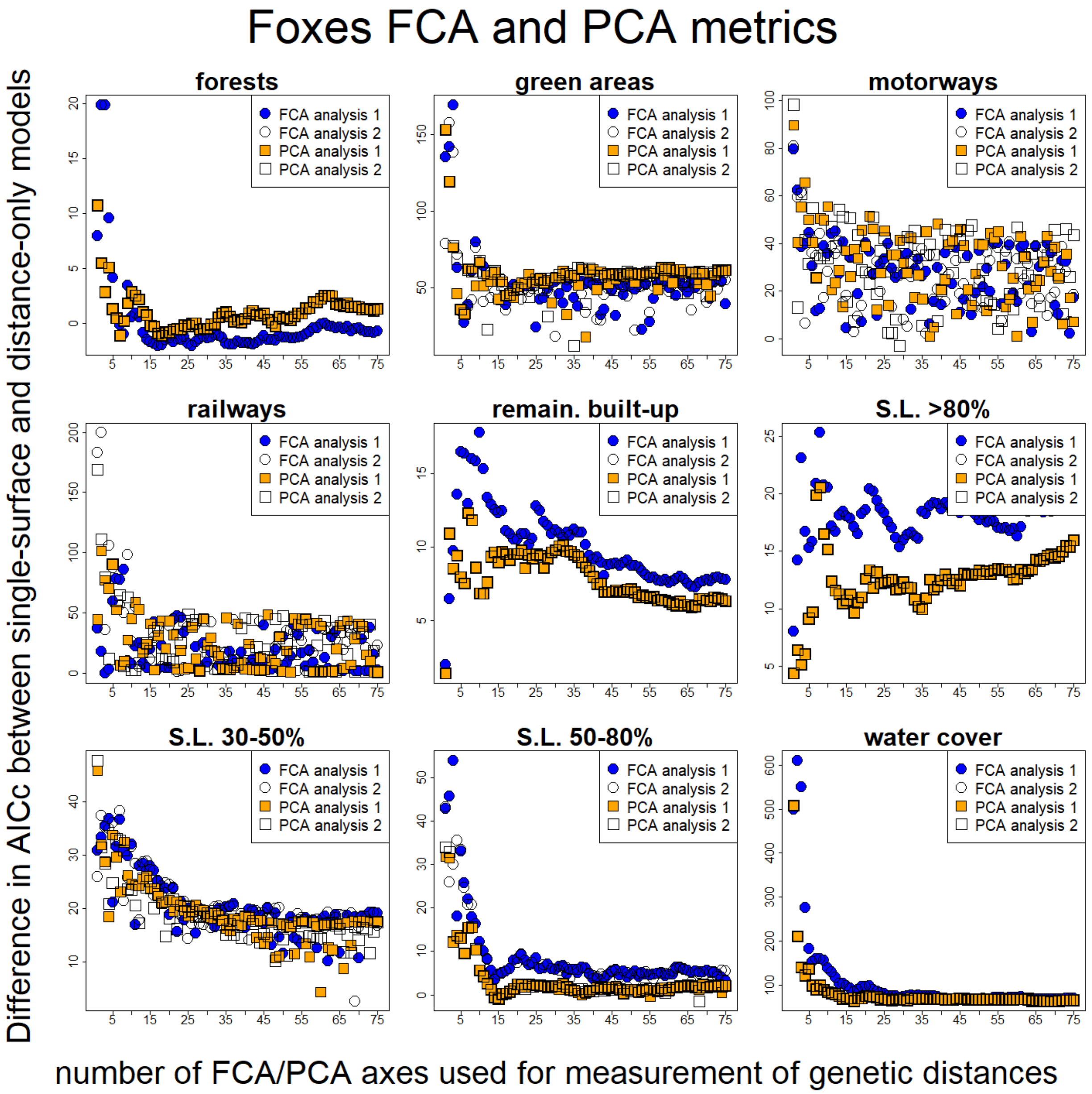

**Figures S7: LIZARDS results of both runs of each landscape factor for all genetic distances metrics derived from FCA/PCA axes 1-75**

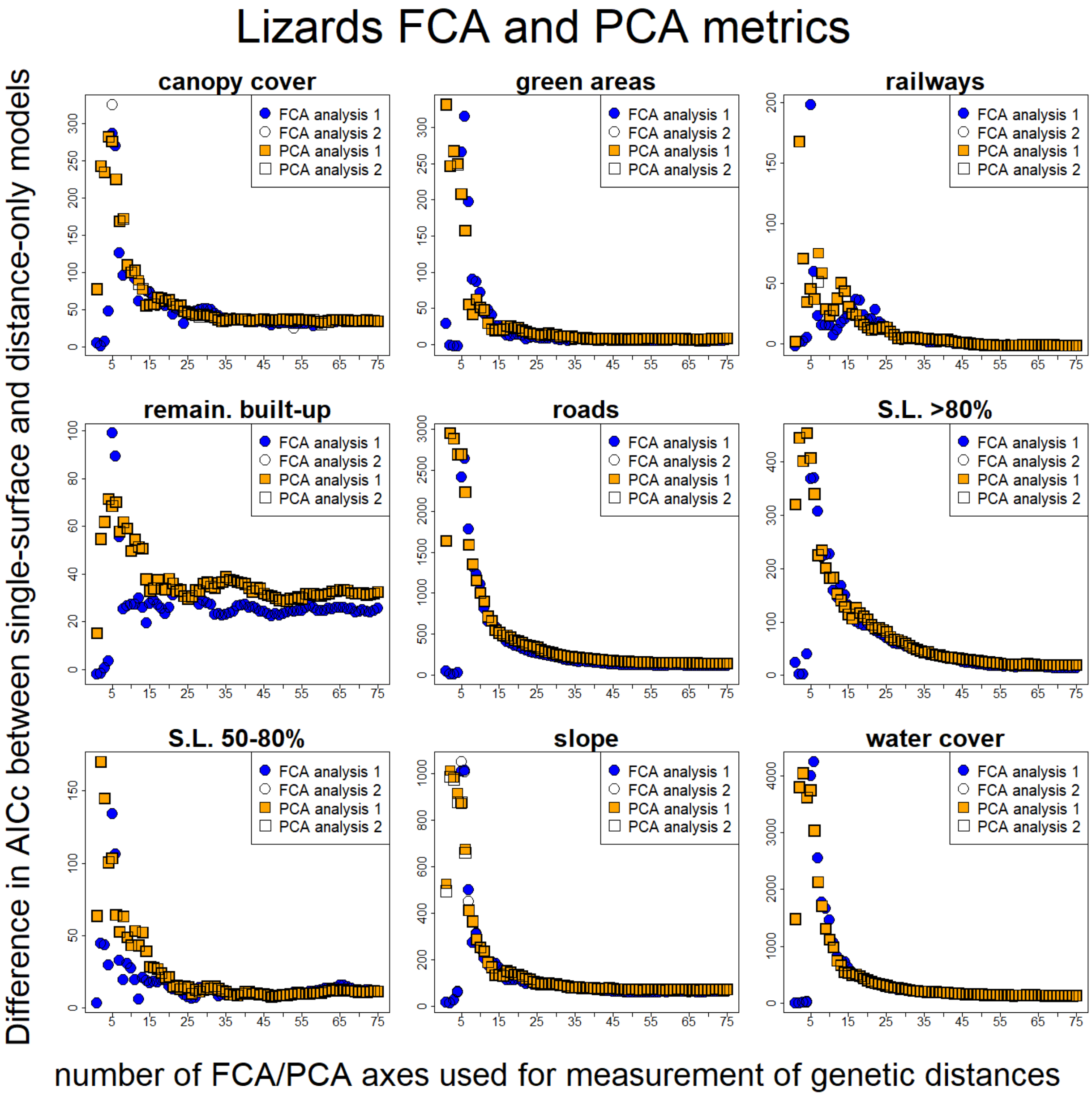

**Figure S8: The differences in Δdist-AICc scores between the initial runs of optimization and the bootstrapping result were mostly small (correlation coefficient = 0.994)**

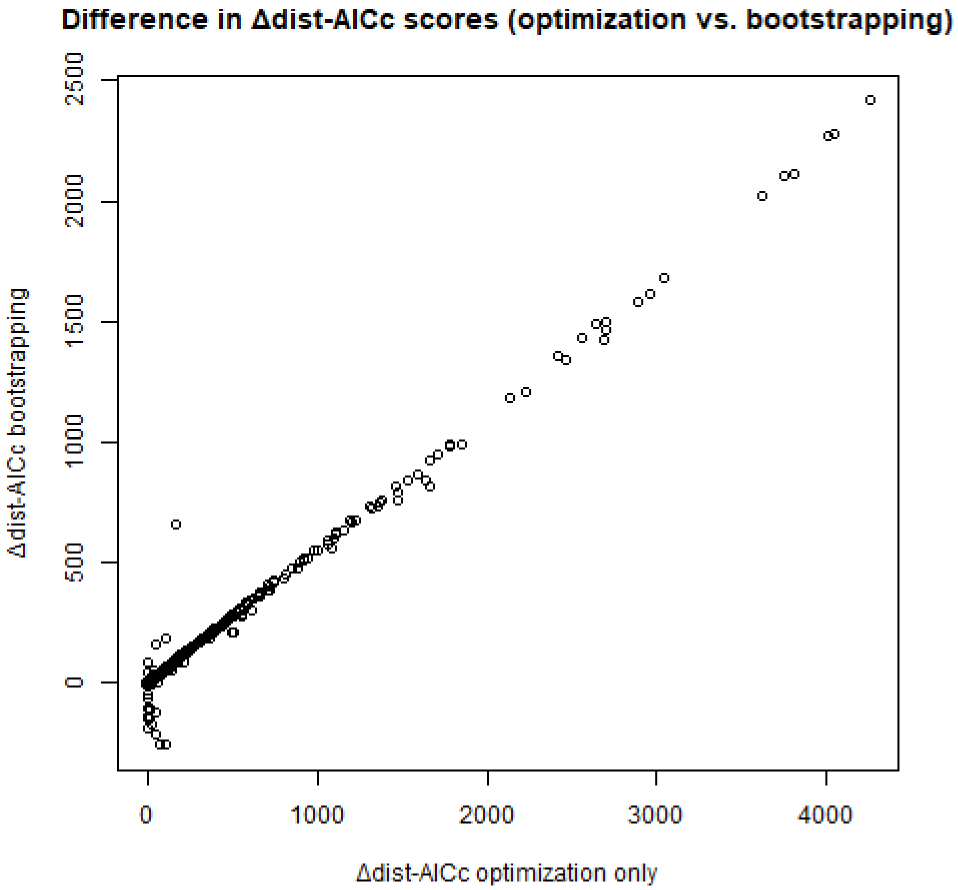

**Figure S9: Difference in Δdist-AICc of models that had a wider parameter space disposable for optimizations compared to models of identical dataset-landscape factor-genetic distance metric combinations that stayed within the set range of resistances (median difference was 4.55 Δdist-AICc.**

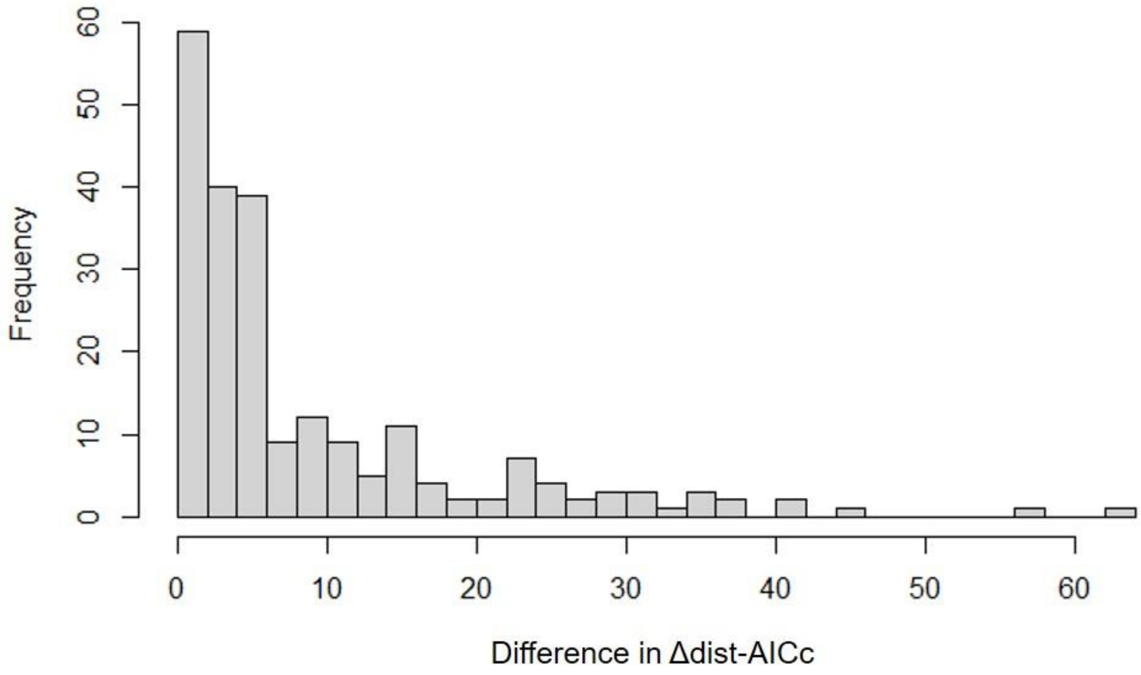

**Figures S10: BOARS results of Δdist-AICc of each landscape factor after bootstrapping for all genetic distances metrics derived from all Other Metrics and FCA/PCA axes 1-10.**

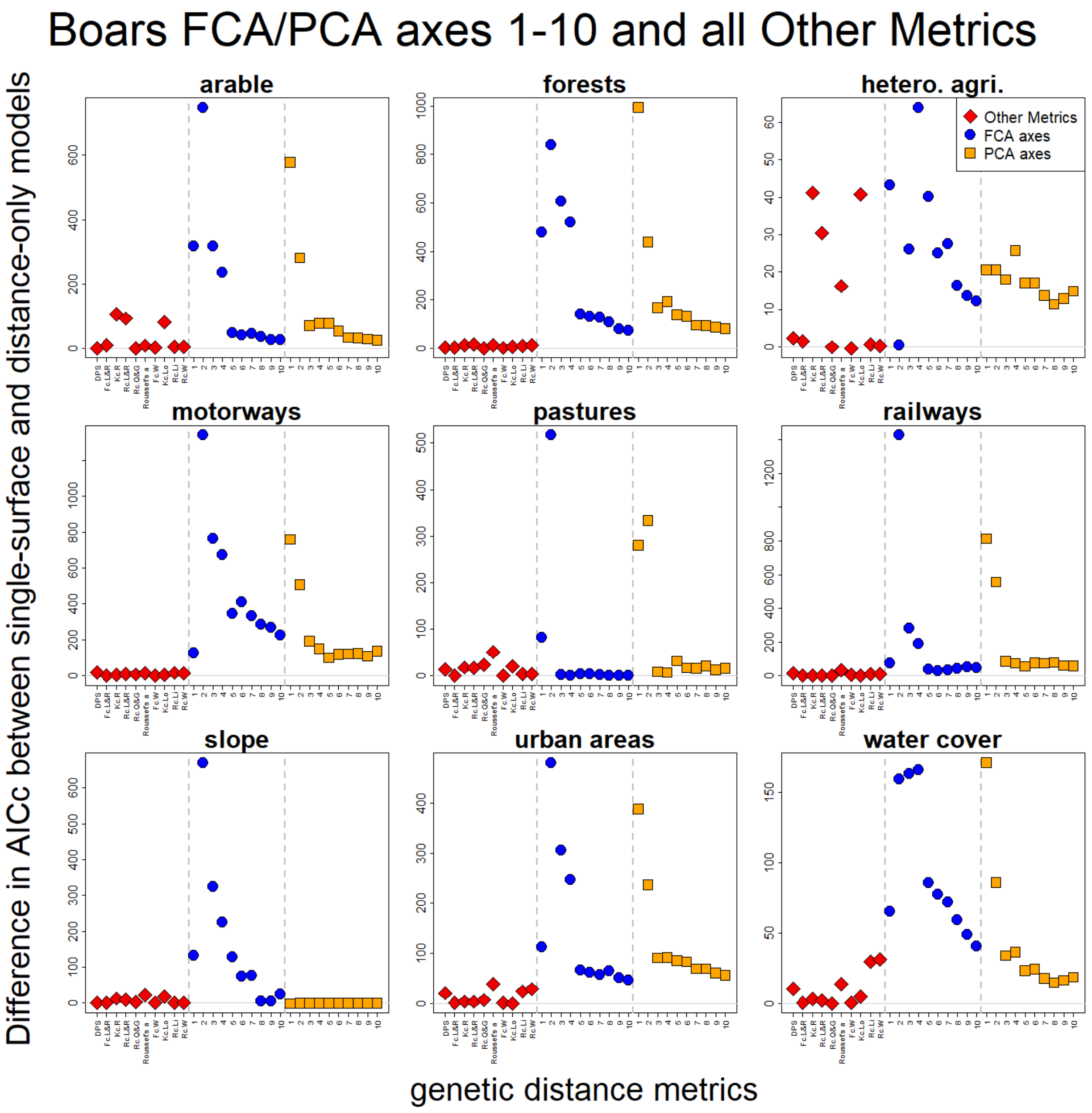

**Figures S11: FOXES results of Δdist-AICc of each landscape factor after bootstrapping for all genetic distances metrics derived from all Other Metrics and FCA/PCA axes 1-10.**

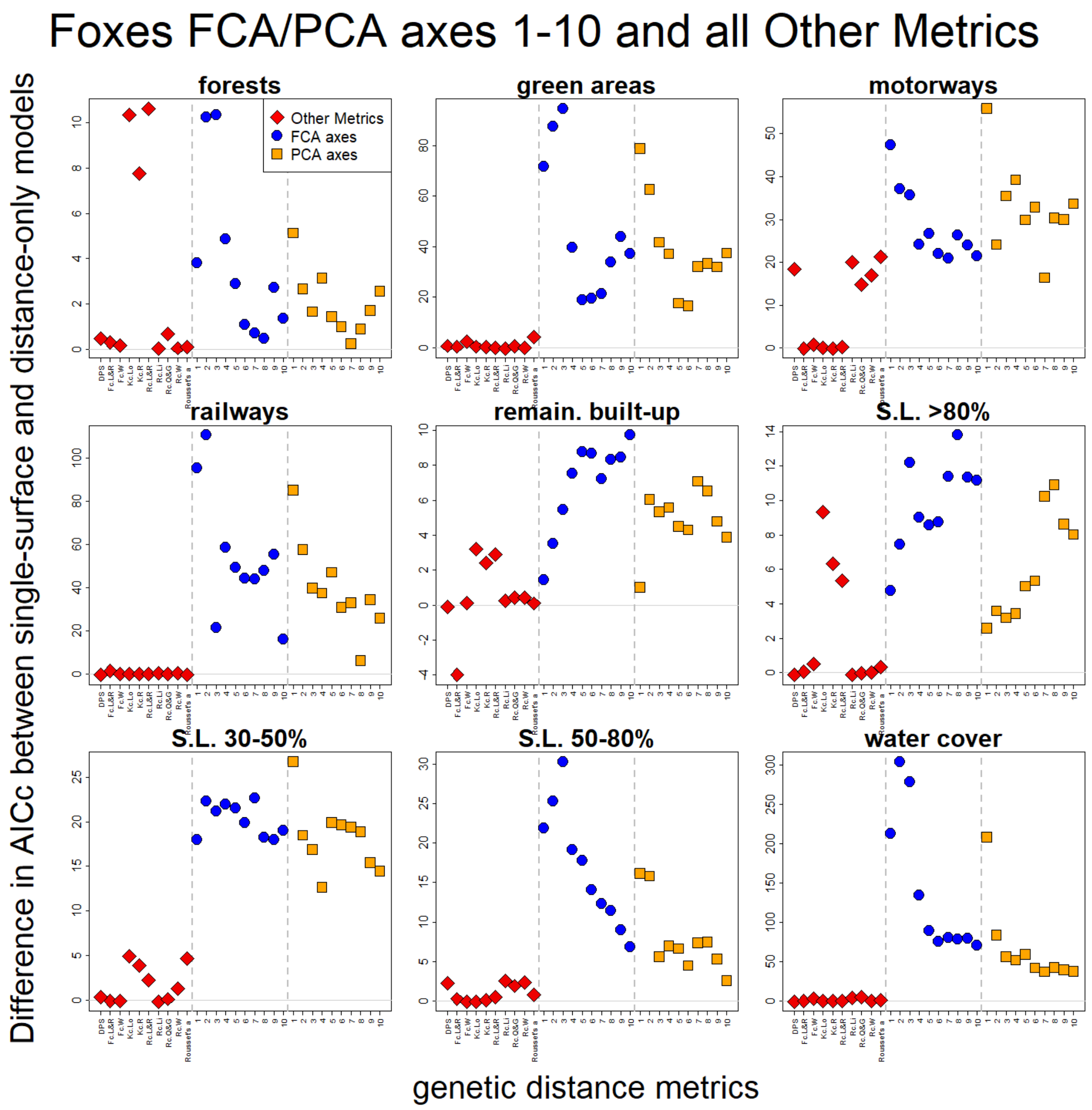

**Figures S12: LIZARDS results of Δdist-AICc of each landscape factor after bootstrapping for all genetic distances metrics derived from all Other Metrics and FCA/PCA axes 1-10.**

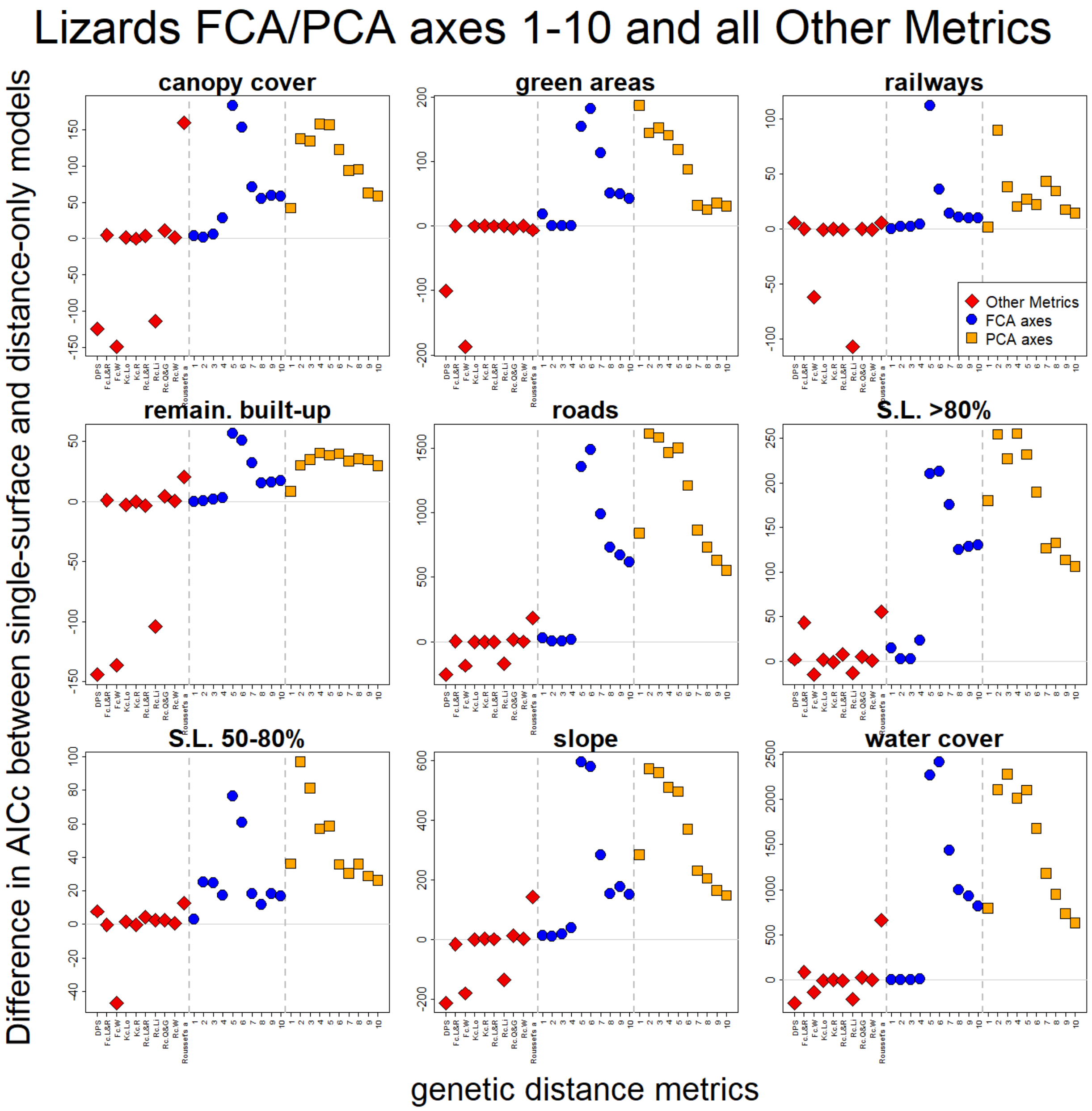

**Figure S13: Resistances of all Other Metrics and the first ten axes of FCA and PCA for BOARS (coloured if model has Δdist-AICc >2).**

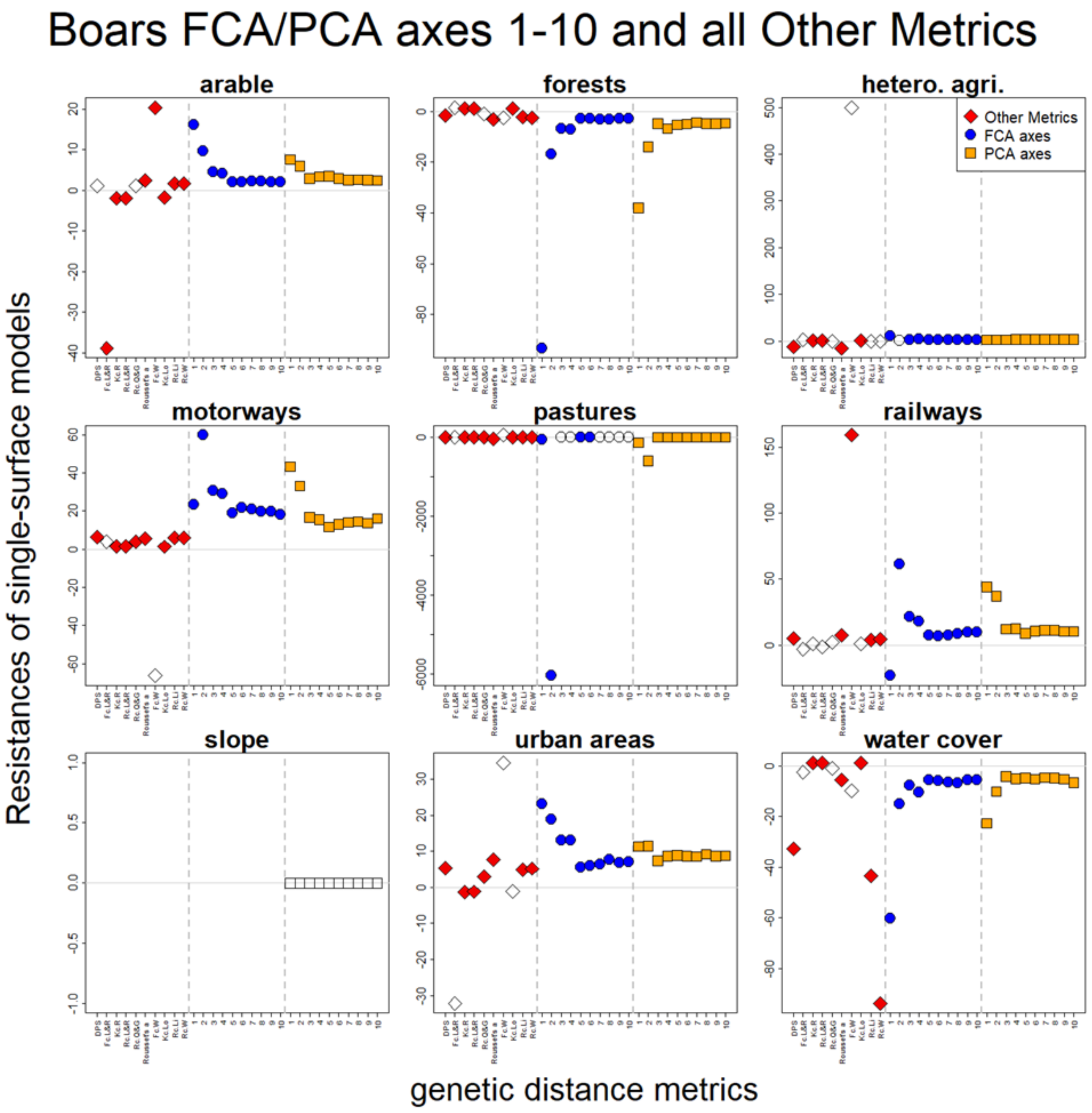

**Figure S14: Resistances of all Other Metrics and the first ten axes of FCA and PCA for FOXES (coloured if model has Δdist-AICc >2).**

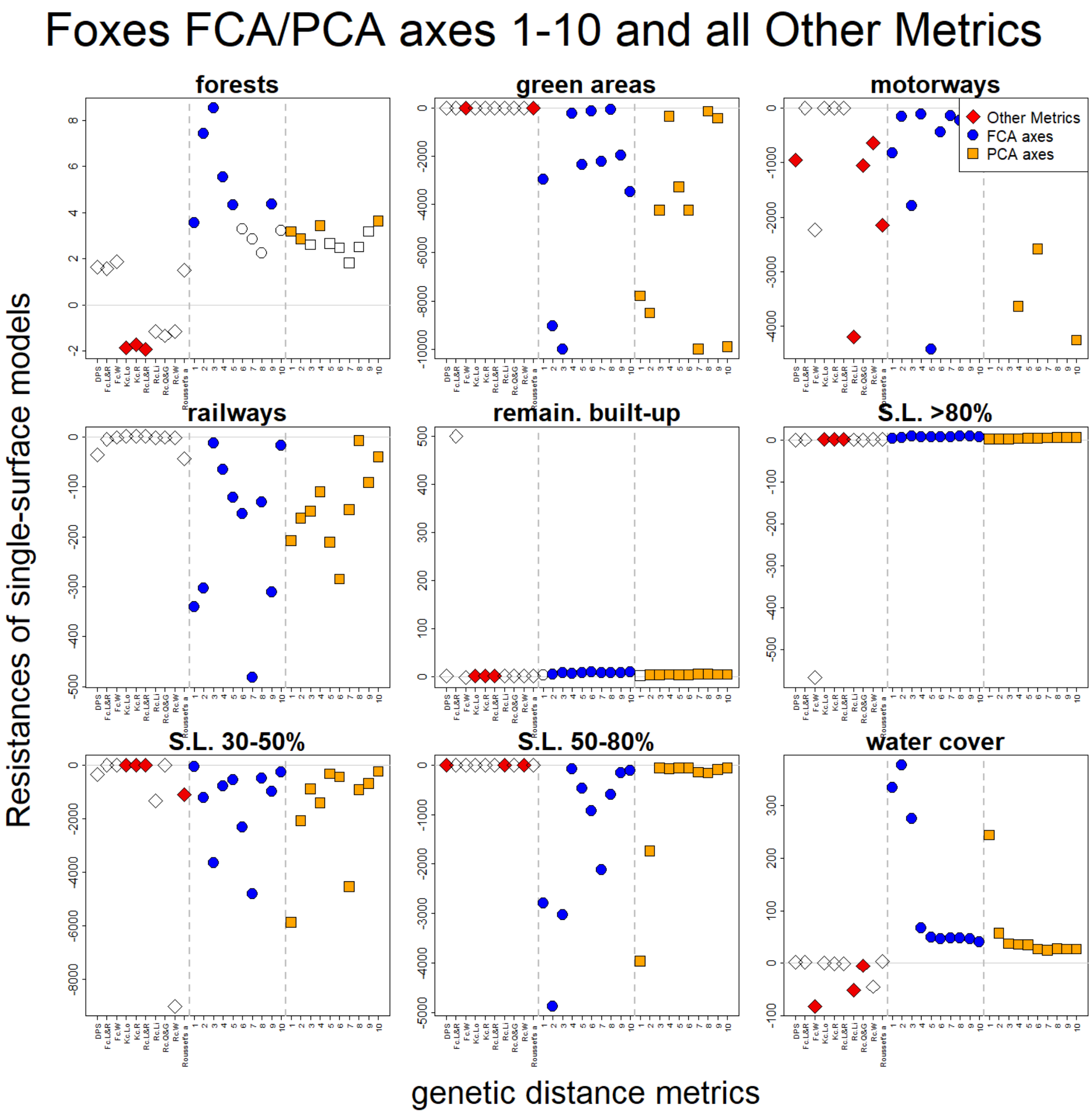

**Figure S15: Resistances of all Other Metrics and the first ten axes of FCA and PCA for LIZARDS (coloured if model has Δdist-AICc >2).**

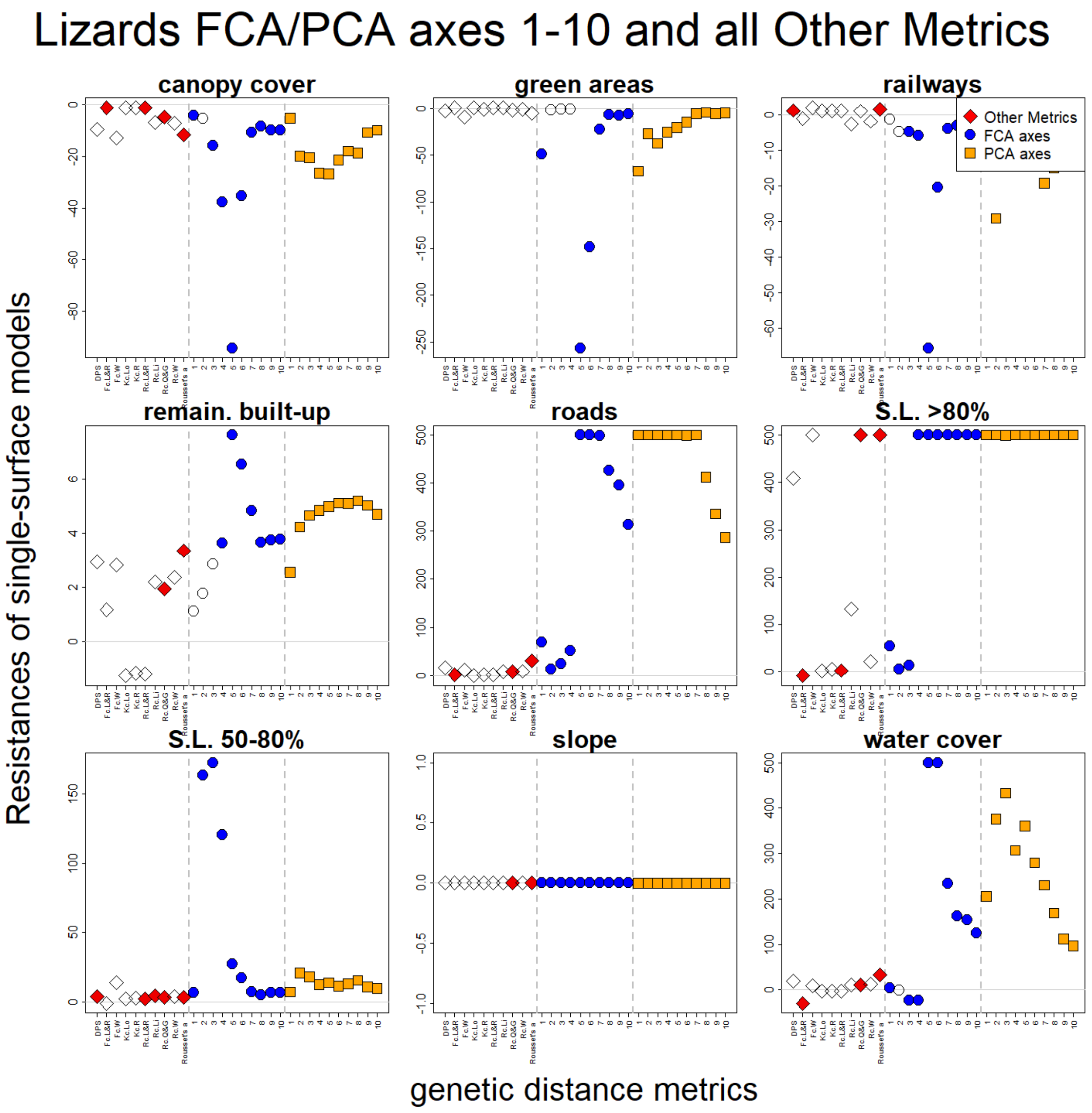
